# Mn^2+^-Induced Structural Flexibility Enhances the Entire Catalytic Cycle and the Cleavage of Mismatches in Prokaryotic Argonaute Proteins

**DOI:** 10.1101/2023.06.28.546964

**Authors:** Lirong Zheng, Bing Zan, Yu Yang, Bingxin Zhou, Banghao Wu, Yan Feng, Qian Liu, Liang Hong

## Abstract

Prokaryotic Argonaute (pAgo) proteins, a class of DNA/RNA-guided programmable endonucleases, have been extensively utilized in nucleic acid biosensors. The specific binding and cleavage of nucleic acids by pAgo proteins, which are crucial processes for their applications, are dependent on the presence of Mn^2+^ bound in the pockets, as verified through X-ray crystallography. However, a comprehensive understanding of how dissociated Mn^2+^ in the solvent affects the catalytic cycle, and its underlying regulatory role in this structure-function relationship, remains underdetermined. By combining experimental and computational methods, this study reveals that unbound Mn^2+^ in solution enhances the flexibility of diverse pAgo proteins. This increase in flexibility through decreasing the number of hydrogen bonds, induced by Mn^2+^, leads to higher affinity for substrates, thus facilitating cleavage. More importantly, Mn^2+^-induced structural flexibility increases the mismatch tolerance between guide-target pairs by increasing the conformational states, thereby enhancing the cleavage of mismatches. Further simulations indicated that the enhanced flexibility in linkers triggers conformational changes in the PAZ domain for recognizing various lengths of nucleic acids. Additionally, Mn^2+^-induced dynamic alterations of the protein cause a conformational shift in the N domain and catalytic sites towards their functional form, resulting in a decreased energy penalty for target release and cleavage. These findings demonstrate that the dynamic conformations of pAgo proteins, resulting from the presence of the unbound Mn^2+^ in solution, significantly promote the catalytic cycle of endonucleases and the tolerance of cleavage to mismatches. This flexibility enhancement mechanism serves as a general strategy employed by Ago proteins from diverse prokaryotes to accomplish their catalytic functions and provide useful information for Ago-based precise molecular diagnostics.

## Introduction

Prokaryotic Argonaute (pAgo) proteins, a class of endonucleases that play a crucial role in DNA interference in prokaryotic organisms (1–3), have gained significant attention in the field of biotechnology and medicine for their ability to target and cleave specific sequences in DNA/RNA (4–11). One of the most important applications is diagnostics, where pAgo proteins can be utilized to design molecular diagnostic assays that detect and quantify specific nucleic acid sequences, such as pathogens or cancer-associated mutations (6, 7, 12–14). These assays can lead to the development of more sensitive and specific diagnostic methods, greatly improving the early detection and accurate treatment of diseases. Another crucial use of pAgo proteins is to label nucleic acids both *in vitro* and *in vivo* (15–18). Their high affinity to the substrate, stability in complex, and specific recognition of the target sequence offer significant advantages over traditional methods that rely on nucleic acid binding, such as DNA-painting and DNA fluorescence in situ hybridization (FISH) technology. Furthermore, the combined action of pAgo proteins and nuclease-deficient RecBC helicase can cleave double-strand DNA (10), potentially leading to the design of gene therapies that specifically target disease-causing genes. These applications of pAgo proteins provide new possibilities for the development of therapeutics and genetic manipulation technologies. pAgo proteins predominantly contain six domains, including the N-terminal domain, Linker1, PIWI-Argonaute-Zwille domain (PAZ), Linker2, middle domain (MID), and P-element Induced Wimpy Testis domain (PIWI) (3, 19, 20). Importantly, the structural dynamics of these domains are critical for the biofunction of pAgo proteins (21–26). The pAgo proteins bind to guide nucleic acids, with the 5’-end and 3’-end of the nucleic acid being accommodated in the basic pocket of the MID domain and the PAZ domain, respectively (27). This guide-loaded complex then searches for complementary target strands through base-pairing with the seed region of the guide strand, which is followed by the propagation of the nucleic acid duplex (21, 27). Subsequently, the 3’-end of the guide is released from the PAZ domain, and the active site of the Glu-finger is closed for correct base-pairing (21). Conformational changes of the catalytic tetrad in the PIWI domain then allow for the cleavage of the target strand (21). Finally, the N domain helps to release the cleaved target, and the protein-guide complex is rearranged to its initial conformation for the next round of cleavage (28, 29).

In particular, the presence of Mn^2+^ is crucial for the catalytic activity of pAgo proteins, as determined by their crystal structures (21, 25, 30–34). Most known pAgo proteins utilize one Mn^2+^ bound to the MID pocket for interactions with the 5’phosphate in the guide strand, and two Mn^2+^ ions bound to the catalytic sites (a DEDX motif, where X is K, D, or H) in the PIWI domain for cleavage of the target strand between the 10^th^ and 11^th^ bases, counted from the 5’-end of the guide (21, 31). Despite the direct role of these bound Mn^2+^, a large number of Mn^2+^ are dissociated and present in the solvent. However, the role of these free ions in the catalytic function of pAgo proteins remains unknown. This situation prompts the question of whether these unbound Mn^2+^ in the solvent affect the functional flexibility of pAgo proteins and, subsequently, influence the endonuclease activity of pAgo proteins.

In this study, we employed a combination of differential scanning fluorimetry (DSF), nano differential scanning calorimetry (nanoDSC), circular dichroism (CD) spectrum synchrotron small-angle X-ray scattering (SAXS), biochemical assays, and molecular dynamics (MD) simulation to investigate the impact of unbound Mn^2+^ on the structural dynamics and endonuclease activity of prokaryotic Argonaute (pAgo) proteins from mesophilic (*Clostridium butyricum* Ago (*Cb*Ago) (31), *Paenibacilllus borealis* Ago (*Pb*Ago) (35), *Pseudooceanicola lipolyticus* Ago (*Pli*Ago) (30), and *Brevibacillus laterosporus* Ago (*Bl*Ago) (35)), thermophilic (*Thermus thermophilus* Ago (TtAgo)) and hyperthermophilic (*Pyrococcus furiosus* Ago (*Pf*Ago) (36), *Methanocaldococcus fervens* Ago (*Mf*Ago) (37) and *Ferroglobus placidus* Ago (*Fp*Ago) (38)) organisms. Our results indicate that the presence of unbound Mn^2+^ significantly reduces the hydrogen bonds in pAgo proteins from mesophiles to hyperthermophiles, while other divalent cations (Mg^2+^, Ca^2+^, and Zn^2+^) do not have such effect. This reduction in hydrogen bonds grants the protein a large degree of structural flexibility, while the overall packing of the protein remains intact. Furthermore, we demonstrated that Mn^2+^-dependent dynamics of pAgo proteins show a higher affinity to substrates and thus greatly promote the binding between the guide and target strands, which is key to facilitating the cleavage activity. Remarkably, Mn^2+^ exerts a significant influence on the precision of target DNA cleavage by the protein-target complex, which increases mismatch tolerance between guide-target pairs by increasing the conformational states. All-atom MD simulations at the conditions mimicking the Mn^2+^-present solvent environment revealed that the flexibility of Link1 and Link2 is greatly enhanced after incubating with Mn^2+^, which induces the conformational change of the PAZ domain to recognize various lengths of guide strands. Additionally, the increased flexibility in the Glue-finger and the PAZ domain could help the propagation of duplexes. Furthermore, the Mn^2+^-induced dynamical change shifts the conformation of catalytic sites and the N domain towards its functional form, lowering the energy penalty for target strand cleavage and release, respectively. Overall, our data revealed that the unbound Mn^2+^-induced flexibility of pAgo proteins is crucial for their catalytic functions, and these findings are generally valid for pAgo proteins obtained from various prokaryotes. Moreover, this study sheds light on the underlying molecular mechanisms of the catalytic cycle of pAgo proteins, providing new avenues for research in the field of prokaryotic immunity systems and precise molecular diagnostics.

## Results

### Mn^2+^ enhances the flexibility of pAgo proteins obtained from mesophiles, thermophiles, and hyperthermophiles

The molecular architecture of long pAgo proteins is characterized by a distinct two-lobed structure, which is a common feature among members of this family (3, 22) (as shown in Figure 1A). The first lobe comprises the N domain, Linker1, and PAZ domain, while the second lobe comprises the Linker2, MID domain, and PIWI domain (as shown in Figure 1B). The phylogenetic position of pAgo proteins studied herein, as well as their structural characteristics, are presented in Figure 1C and Figure S1, respectively. Although the overall packing topology of pAgo proteins obtained from different prokaryotes exhibits a similar tertiary and secondary structure (as shown in Figure S1), their functions and optimal physiological temperatures are vastly divergent (3, 24).

**Figure 1.**
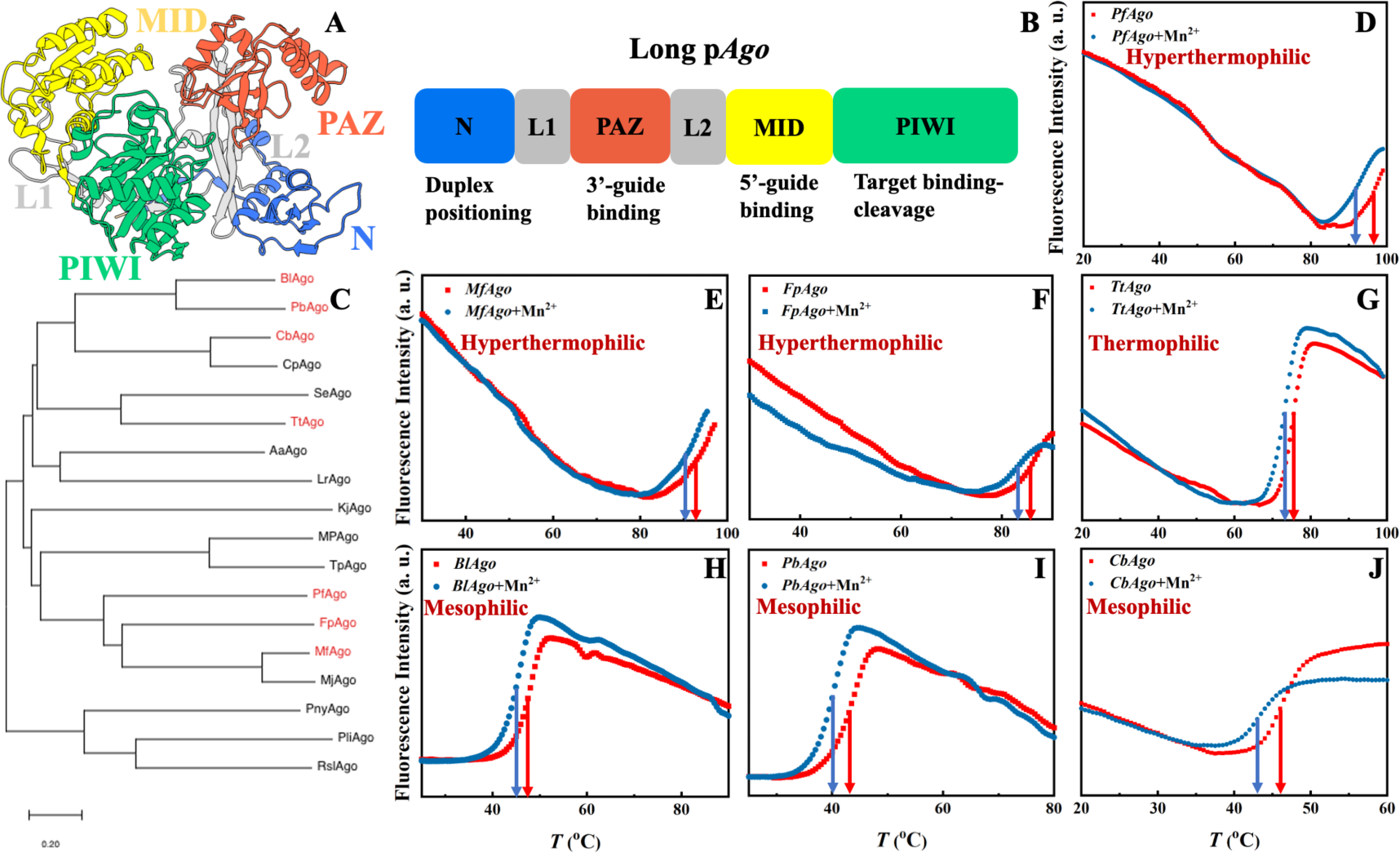
Structural analysis of hyperthermophilic, thermophilic, and mesophilic pAgo proteins incubated with and without Mn^2+^ at different temperatures. (A) Representation of the domain architectures of long pAgo proteins. PAZ domain, MID domain, N domain, and PIWI domain are colored in red, yellow, blue, and green, respectively. (B) Schematic diagram of the domain organization of long pAgo proteins. (C) Maximum likelihood phylogenetic tree of characterized pAgo proteins. DSF spectra of (D) *Pf*Ago, (E) *Mf*Ago, (F) *Fp*Ago, (G) *Tt*Ago, (H) *Bl*Ago, (I) *Pb*Ago, and (J) *Cb*Ago incubated with and without Mn^2+^ measured at different temperatures. The concentration of Mn^2+^ is 5 mM, which is widely applied in the biochemical assay of pAgo proteins. The solid arrows in panels D-J indicate the melting temperature, *T*_m_.

We first examined how the secondary structures of pAgo proteins vary with the concentration of Mn^2+^ at their respective physiological temperatures. The study was conducted using CD spectroscopy, a technique that measures the secondary structure of a protein by analyzing its absorption of circularly polarized light (39). The results, as shown in Figures S2A-C, indicate that the presence of Mn^2+^ does not significantly alter the secondary structures of pAgo proteins. It should be noted that we use Mn^2+^ in this study rather than Mg^2+^ because *Pf*Ago cannot perform its functions when using Mg^2+^ (36).

We then characterized the effect of Mn^2+^ on the thermostability of pAgo proteins by using DSF spectroscopy. The DSF method is a quantitative method that allows for the assessment of thermostability by measuring changes in fluorescence intensity as the temperature is progressively increased, which monitors the tertiary structural changes in proteins (40). The melting temperature (*T*_m_) of the tertiary structure of different pAgo proteins was determined with and without the presence of Mn^2+^. As shown in Figures 1D-J, the results revealed that the addition of Mn^2+^ resulted in a significant decrease in the *T*_m_ of pAgo proteins, reducing the thermostability, whereas other divalent cations (Mg^2+^, Ca^2+^, and Zn^2+^) do not have this effect (Figures S2D-E). To further validate these findings, the melting temperature of pAgo proteins was also determined using nanoDSC, a technique that measures the heat capacity of a protein as a function of temperatures (41). The results obtained through nanoDSC were consistent with those obtained through DSF spectroscopy, showing a similar trend of increased denaturation in the presence of Mn^2+^ (as shown in Figure S3). It is noteworthy that no significant difference was observed when the *T*_m_ of the secondary structure of pAgo proteins was evaluated under similar conditions (see Figure S4). Thus, the above results suggest that the presence of Mn^2+^ may disrupt interactions between amino acids participating in the tertiary structure of the protein but not those involved in the secondary structure (alpha-helix and beta-sheet), thereby reducing the thermostability of the protein.

In order to gain a deeper understanding of the structural factors that contribute to the observed difference in *T*_m_ between pAgo proteins incubated with Mn^2+^ and those incubated without, we conducted all-atom molecular dynamics (MD) simulations on mesophilic (*Cb*Ago), thermophilic (*Tt*Ago), and hyperthermophilic (*Fp*Ago) pAgo proteins incubated with and without Mn^2+^. The example of snapshots of MD simulations is shown in Figure S5 (The example of the potential energy of protein as a function of MD simulation time is shown in Figure S6). Our simulations reveal that the pAgo proteins incubated without Mn^2+^ exhibit a greater number of hydrogen bonds when compared to those incubated with Mn^2+^, while the number of salt bridges remains constant (see Table 1). This disparity in intermolecular interactions, i.e., loss of hydrogen bonds, could reduce the energy barrier for protein unfolding in pAgo proteins incubated with Mn^2+^, which explains the decreased *T*_m_ observed in pAgo proteins observed in Figures 1D-J.

**Table 1.** Number of hydrogen bonds and salt bridges of pAgo proteins incubated with and without Mn^2+^.

| Protein | *FpAgo* | *FpAgo*+$Mn^{2+}$ | *TtAgo* | *TtAgo*+$Mn^{2+}$ | *CbAgo* | *CbAgo*+$Mn^{2+}$ |
| --- | --- | --- | --- | --- | --- | --- |
| Hydrogen bond* | 611 | 588 | 452 | 438 | 307 | 282 |
| Salt bridge* | 13 | 13 | 11 | 11 | 9 | 9 |

\*Number of hydrogen bonds and salt bridges are averaged over the entire simulation time.

By integrating the results of CD, DSF, and MD simulations, it suggests that Mn^2+^ possesses the capacity to perturb the intermolecular interactions among amino acid residues participating in the building of the tertiary structure, while not for the secondary structure (see Table S1).

It has been reported that structural flexibility is crucial for the biofunction of pAgo proteins (22, 24). The intriguing question then arises as to whether the decreased intermolecular interactions resulting from the addition of Mn^2+^ can increase the flexibility of pAgo proteins. To address this, we conducted a structural analysis of four different types of pAgo proteins, namely *Fp*Ago (hyperthermophilic), *Tt*Ago (thermophilic), *Bl*Ago (mesophilic), and *Cb*Ago (mesophilic) using synchrotron small-angle X-ray scattering (SAXS) and Porod-Debye analysis (24, 42, 43). The Porod-Debye analysis, which is a combination of the scattering intensity and the fourth power law of the scattering wavevector, *q*^4^·*I*(*q*), can be utilized to reveal the flexibility of biomacromolecules (42). An unambiguous Porod plateau in the scattering profile indicates the protein takes a rigid structure, while a lack of a plateau implies that the biomacromolecule forms dynamic conformations (42). As shown in Figures 2A-D, the Porod-Debye plots of *Fp*Ago, *Tt*Ago, *Bl*Ago, and *Cb*Ago all displayed a clear Porod plateau. However, after incubation with Mn^2+^, a significant change in the Porod-Debye region was observed, with a loss of the plateau suggesting the protein flexibility is greatly increased in the presence of Mn^2+^ (see Figures 2A-D). To further support these findings, we compared the flexibility of pAgo proteins incubated with and without Mn^2+^ using root mean square fluctuation (RMSF) derived from MD simulations. The RMSF of pAgo proteins incubated with Mn^2+^ is found to be larger than pAgo proteins incubated without Mn^2+^, indicating that Mn^2+^ confers increased flexibility to pAgo proteins (Figures 2E-H). Our findings demonstrate that Mn^2+^ has a universal effect to enhance the structural dynamics of pAgo proteins from different prokaryotes.

**Figure 2.**
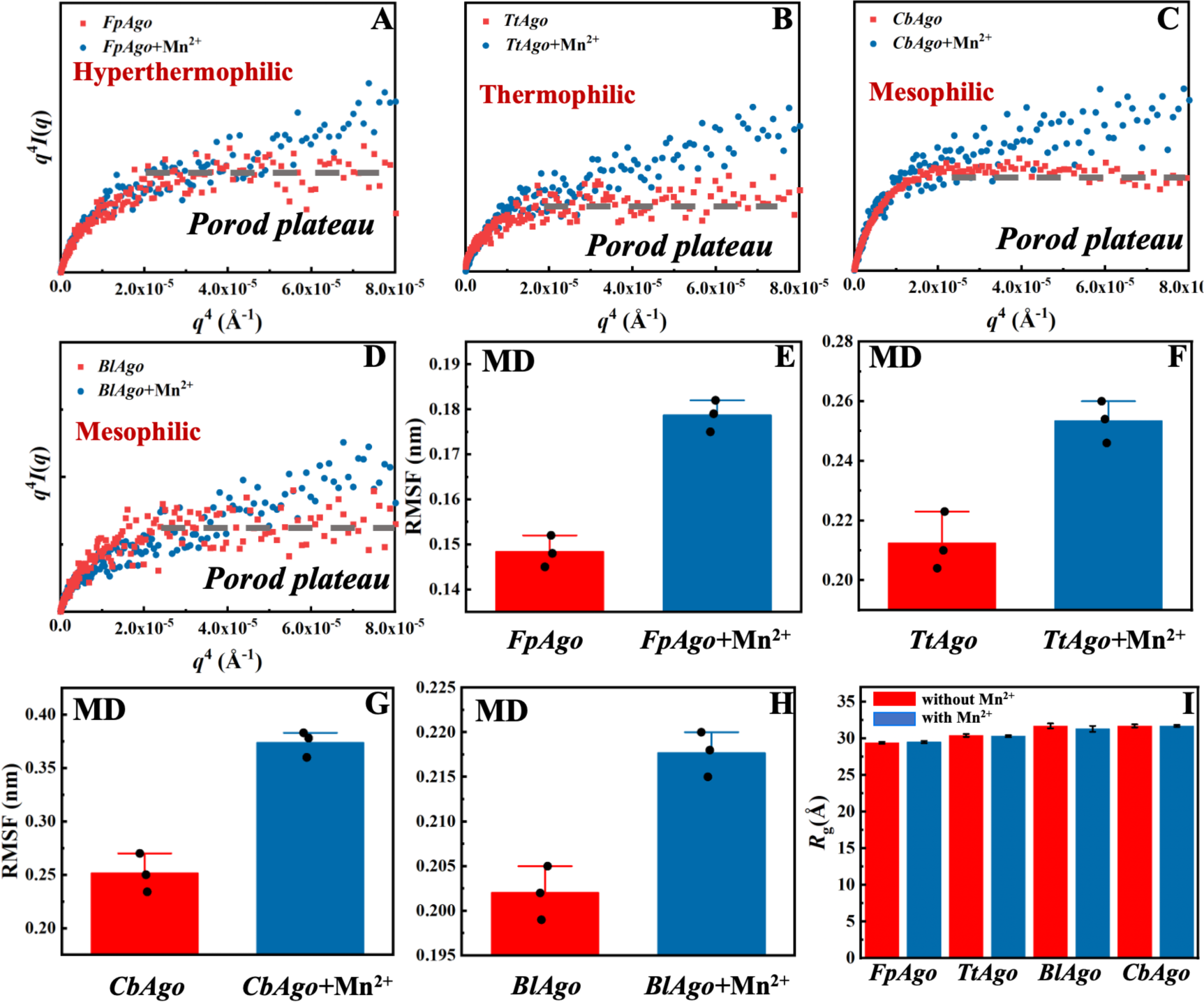
Structures and dynamics of pAgo proteins incubated with and without Mn^2+^ derived from SAXS and MD simulation. Porod-Debye analysis, *q^4^I*(*q*) vs. *q^4^*, of (A) *Fp*Ago, (B) *Tt*Ago, (C) *Bl*Ago, and (D) *Cb*Ago incubated with and without Mn^2+^. The concentration of Mn^2+^ is 5 mM. The dashed gray line in each panel indicates the Porod plateau. The details about the SAXS data collection and analysis are summarized in Materials and Methods. Comparison of root mean squared fluctuation (RMSF) of (E) *Fp*Ago, (F) *Tt*Ago, (G) *Cb*Ago, and (H) *Bl*Ago incubated with and without Mn^2+^. RMSF of pAgo proteins is averaged over entire residues. The results from three independent MD simulations were quantified. Error bars represent the standard deviations of three independent simulations. The detailed procedure is presented in the Materials and Methods. (I) The radius of gyration (*R*_g_) of *Fp*Ago, *Tt*Ago, *Bl*Ago, and *Cb*Ago incubated with and without Mn^2+^ derived from SAXS. The Guinier plots of proteins are shown in Figures S7-S17.

Furthermore, we also compared the structural compactness of pAgo proteins incubated with and without Mn^2+^. By utilizing Guinier analysis (44, 45), one can derive the radius of gyration (*R*_g_) of the protein molecule, and we found that *R*_g_ of pAgo proteins incubated with Mn^2+^ is comparable to that of pAgo proteins incubated without Mn^2+^ (Figures 2I, S7-S17, and Table S3), which is in agreement with the *R*_g_ derived from MD simulations (Figure S18). Furthermore, the Porod volume of pAgo proteins incubated with and without Mn^2+^ was found to be similar, indicating that Mn^2+^ does not have a significant impact on the overall compactness of pAgo proteins. These findings were also supported by the *ab initio* low-resolution model reconstructed from SAXS data. As shown in Figure S19, the *ab initio* structure of pAgo proteins incubated with Mn^2+^ is found to be similar to that of pAgo proteins incubated without Mn^2+^.

The findings from DSF, CD, nanoDSC, SAXS, and MD simulation studies provide a convincing explanation of Mn^2+^’s impact on the structural flexibility of pAgo proteins, which suggests that the presence of Mn^2+^ greatly enhances the protein flexibility while preserving the protein’s secondary structure. The observation that the overall structure of pAgo proteins remains similar to the crystalline form when incubated with Mn^2+^ further supports this conclusion, indicating that the flexibility changes are not due to significant alterations in the protein’s overall structure.

### Enzymatic functional role of the Mn^2+^ induced flexibility in pAgo proteins

The physical characterization of Mn^2+^ has revealed that it can enhance the flexibility of pAgo proteins. However, the implications of this increase in flexibility on the catalytic function of pAgo proteins remain unclear. As shown in Figure 3A, the catalytic cycle of pAgo proteins involves several functional steps, including binding guide (gDNA binding), target recognition and annealing (tDNA binding), target cleavage, and release (tDNA cleaving and releasing) (22, 46). In order to study the impact of increased protein structural flexibility on guide DNA binding, we employed the fluorescence polarization assay on *Pli*Ago (DSF measurements of *Pli*Ago incubated with and without Mn^2+^ are shown in Figure S20A), a unique protein that does not require Mn^2+^ for substrate binding (30). The results, as shown in Figure 3B, indicate that as the concentration of Mn^2+^ increases, the dissociate constant *K*_d_ decreases, thus demonstrating an increased binding of gDNA to the protein and the formation of the more protein-gDNA binary complex. To further verify these results, we conducted gDNA binding experiments on *Cb*Ago using a mutation on the binding site amino acids, resulting in a binding pocket in the MID domain that does not bind Mn^2+^ (the amino acids sequence is shown in Table S5) (30). We found that the *Cb*Ago mutant (*Cb*Ago*M*) incubated with Mn^2+^ had a greater binding affinity to gDNA than it incubated without Mn^2+^ (as shown in Figure 3C), providing additional support for the notion that the increase in protein flexibility caused by Mn^2+^ can enhance the binding ability of proteins to gDNA (DSF measurements of *Cb*Ago*M* incubated with and without Mn^2+^ are shown in Figure S20B). Additionally, an increased quantity of protein-gDNA binary complexes can effectively function as templates, thereby facilitating the binding of a greater amount of tDNA.

**Figure 3.**
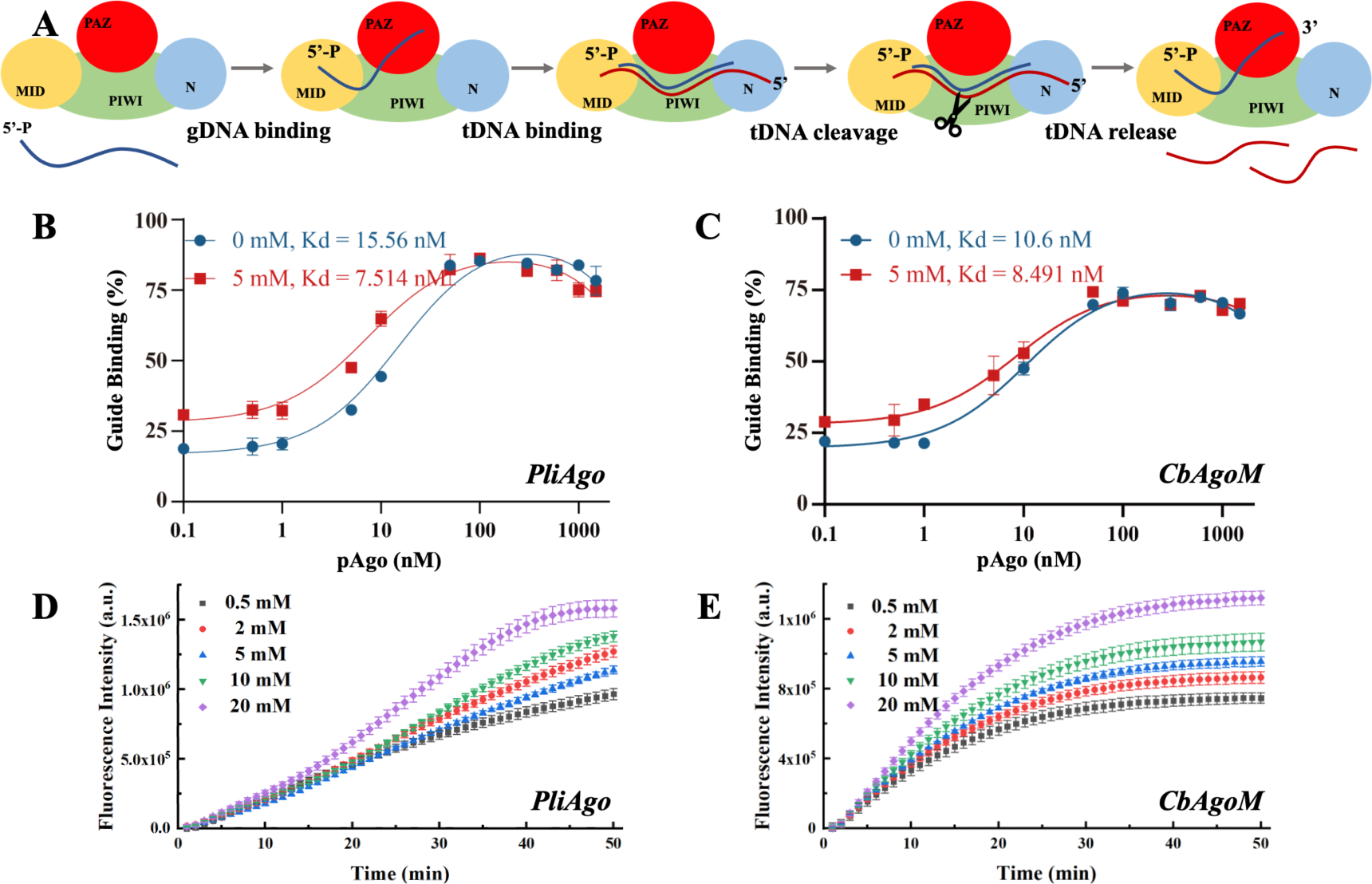
The role of the Mn^2+^-dependent structural dynamics of pAgo proteins for binding guide DNA and cleavage of target DNA. (A) Schematic diagram of DNA-catalytic cycle of pAgo proteins. Fluorescence polarization assay of the binding of (B) *Pli*Ago and (C) *Cb*Ago*M* to gDNA at different concentrations of Mn^2+^. Cleavage assays of (D) *Pli*Ago and (E) *Cb*Ago*M* incubated with different concentrations of Mn^2+^. The cleavage activity of pAgo proteins is traced by fluorescence intensity as a function of time. In all experiments, protein, guide, and target were mixed at a 5:1:1 molar ratio and incubated at 37 °C. The reaction buffer without Mn^2+^ contains 10 mM EDTA. Three samples were used for each experimental condition. The results from three independent experiments were quantified. Error bars represent the standard deviations of three independent experiments. The detailed procedure is presented in the Materials and Methods. The nucleotide sequences of the gDNA and tDNA are presented in Table S4.

The process of cleavage and release of tDNA is of paramount importance for the subsequent rounds of tDNA cleavage mediated by pAgo proteins (22). In order to investigate the impact of Mn^2+^ on the cleavage efficiency of pAgo proteins on tDNA, we employed a highly sensitive fluorescence-based assay. To this end, we labeled the 5’- and 3’-end of tDNA with the fluorescent group 6’FAM and quenching group BHQ1, respectively. This allowed us to monitor the cleavage and release process of pAgo proteins on tDNA in real-time, with the absence of a fluorescent signal indicating that tDNA was either not cleaved or had been cleaved but not yet released, and the presence of a fluorescent signal indicating that tDNA had been cleaved and released successfully. As depicted in Figures 3D and E, the cleavage rate and yield of cleavage products of *Pli*Ago and *Cb*Ago*M* on tDNA exhibited a clear dependence on the concentration of Mn^2+^, with tested higher concentrations resulting in increased cleavage products. Our findings provide important insights into the role of Mn^2+^ in modulating the catalytic activity of pAgo proteins, indicating that the enhancement of protein flexibility brought about by Mn^2+^ might facilitate conformational changes during the cleavage process and the release of products.

It has been reported that small interfering DNA of varying lengths present inside cells can function as gDNA for proteins, which subsequently cleave invading DNA (47). In light of this, we explored the flexibility of pAgo proteins and their potential roles in facilitating binding to tDNA and gDNA of different lengths. Our analysis, illustrated in Figure S21, revealed that when the concentration of Mn^2+^ increases from 0.1 mM to 5 mM, *K*_d_ for the protein with tDNA and gDNA of various lengths decrease, indicating enhanced binding of gDNA and tDNA to the pAgo protein. These findings suggest that the ability of pAgo proteins to adapt their conformation, facilitated by an increase in structural flexibility induced by Mn^2+^, enables them to bind to tDNA and gDNA of varying lengths.

We then investigated the effect of Mn^2+^-induced flexibility in proteins on the cleavage and release of tDNA of varying lengths. Our experiments utilized a range of Mn^2+^ concentrations to measure the rate of tDNA cleavage and release. As shown in Figure S22, the results revealed a statistically significant positive correlation between increasing concentrations of Mn^2+^ and the fluorescence intensity per unit of time. This observation demonstrates an augmentation in the rate of tDNA cleavage and release in response to Mn^2+^-induced changes in pAgo protein flexibility, thus suggesting that the enhancement of pAgo protein flexibility serves as a facilitative mechanism for the cleavage and release of tDNA.

### Mn^2+^-dependent precision cleavage of target DNA

It’s widely recognized that the efficiency of pAgo’s cleavage is influenced by mismatches that occur between the guide and the target, particularly within the seed region. The presence of these mismatches can cause “bubbles” to form between the guide and the target, consequently affecting their mutual recognition and the subsequent cleavage by pAgo (Figure 4A). In this section, we also mutated the amino acids in the binding pocket of pAgo proteins to exclude the impact of Mn^2+^ on binding gDNA and tDNA (30). In the case of *Pf*Ago, the introduction of a single nucleotide mismatch at varying positions within the guide can cause weakening or even elimination of target cleavage (Figure 4B). Interestingly, when the concentration of Mn^2+^ is increased, *Pf*Ago can cleave the target even with a mismatched guide (Figure 4B). This suggests that Mn^2+^ enhances the flexibility of *Pf*Ago, thereby increasing its tolerance for mismatches. We then selected three guides that either perfectly matched the target sequence (guide-m0) or contained mismatches at the fourth (guide-m4) and seventh (guide-m7) positions. We performed cleavage assays at gradually increasing Mn^2+^ concentrations (Figures 4C and D). Under conditions of 0.1 mM Mn^2+^, the three guides directed varied catalytic efficiency, ranging from 20% to 70%. Conversely, when the Mn^2+^ concentration was increased to 1 mM, the cleavage efficiency of all three guides reached a comparable level. Similarly, *Cb*Ago demonstrated enhanced activity with increasing Mn^2+^ concentrations, as shown in Figure S23.

**Figure 4.**
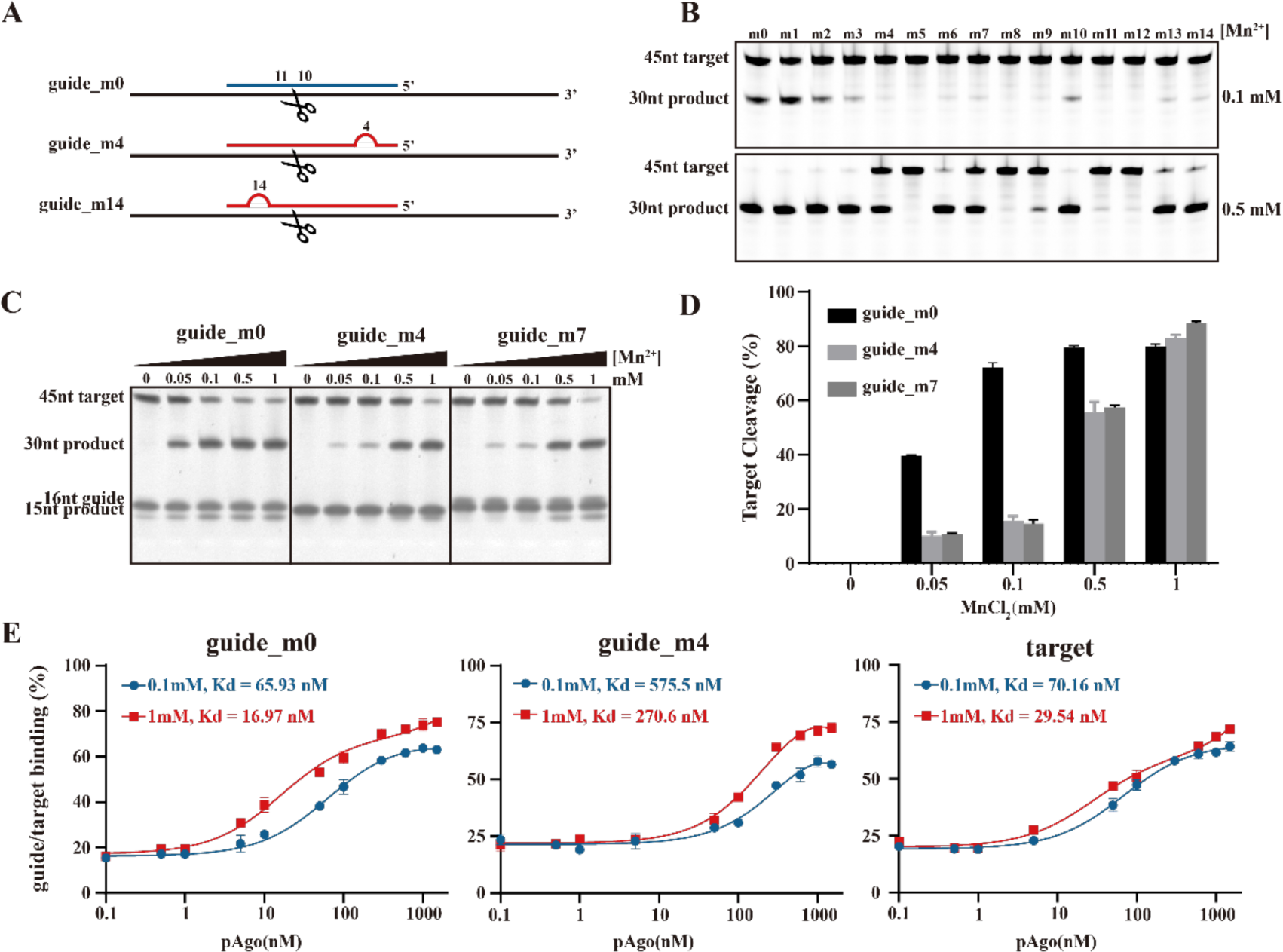
The role of the Mn^2+^-dependent structural dynamics of pAgo proteins for the designed mismatched guide DNA directed cleavage on target DNA. (A) Schematic representation of the designed mismatched gDNA directed cleavage on tDNA. The introduced mismatched nucleotides are indicated by the red semicircle. The nucleic acids sequence of gDNA and tDNA are shown in Table S6. *Pf*Ago is used here. (B) and (C) Gel detection results for the designed mismatched gDNA-directed cleavage on tDNA at different concentrations of Mn^2+^. (D) Quantifying the effects of incubating with Mn^2+^ on tDNA cleavage obtained from panel (C). (E) Fluorescence polarization assay of the designed mismatched guide and target at different concentrations of Mn^2+^. The results from three independent experiments were quantified. Error bars represent the standard deviations of three independent experiments. It should be noted that we mutated the amino acids in the binding pocket of pAgo proteins to exclude the impact of Mn^2+^ on binding gDNA and tDNA.

In order to further investigate the role of Mn^2+^, we conducted fluorescence polarization experiments to measure the apparent dissociation constants (*K*_d_) of *Pf*Ago with either the guide or the target at different Mn^2+^ concentrations. We observed that an increase in the concentration of Mn^2+^ enhances the affinity of *Pf*Ago for both the guide and the target. For the guides, the *K*_d_ values increased up to 4-fold and 2-fold for guide-m0 and guide-m4, respectively. For the target, the *K*_d_ values increased 3-fold (Figure 4E). These findings suggest an increased binding affinity of the protein for mismatched gDNA and tDNA, which attributes to increased protein flexibility under elevated Mn^2+^ concentrations. This provides further support for our hypothesis that Mn^2+^ may play a crucial role in enhancing the tolerance of DNA cleavage to mismatches.

Thus, the data obtained indicate that the enhancement of flexibility in pAgo proteins, induced by Mn^2+^, broadens the tolerance for mismatches between the gDNA and the tDNA. This heightened tolerance, however, incurs a trade-off, leading to a consequent decrease in the precision of tDNA cleavage orchestrated by the protein. This pivotal finding underscores the profound implications of the interplay between protein flexibility and nucleic acid detection accuracy, highlighting the necessity for maintaining an optimal balance between these two critical factors.

### Mn^2+^-dependent dynamical states in pAgo protein derived from MD simulations

To gain a more comprehensive understanding of the effects of Mn^2+^ on the conformational dynamics related to the function of pAgo proteins at the atomic scale, we conducted a detailed examination using MD simulations. This allowed us to analyze structural changes and flexibility in response to the presence of Mn^2+^. Figure 5A compares the structure of *Cb*Ago incubated with Mn^2+^ (represented in blue) to that of *Cb*Ago without Mn^2+^ (represented in red) over trajectory time. The results show that Mn^2+^ leads to a significant increase in the conformational dynamics of *Cb*Ago compared to it incubated without Mn^2+^, indicating that Mn^2+^ notably enhances the flexibility of the proteins (as shown in Figure 5B). However, the content of the alpha helix, beta sheet, and turns and loops as terms of secondary structure, as well as the radius of gyration, remain consistent regardless of the presence of Mn^2+^, in line with the SAXS and CD results presented in Figures 1 and 2. Additionally, when comparing the conformational phase space of *Cb*Ago sampled in the MD simulation with and without Mn^2+^, we found that the presence of Mn^2+^ allows the protein to explore a significantly larger conformational space (as depicted in Figures 5C and D). This larger conformational space suggests that the presence of Mn^2+^ may lower the energy barrier between different states, thus facilitating the transition between functional states.

**Figure 5.**
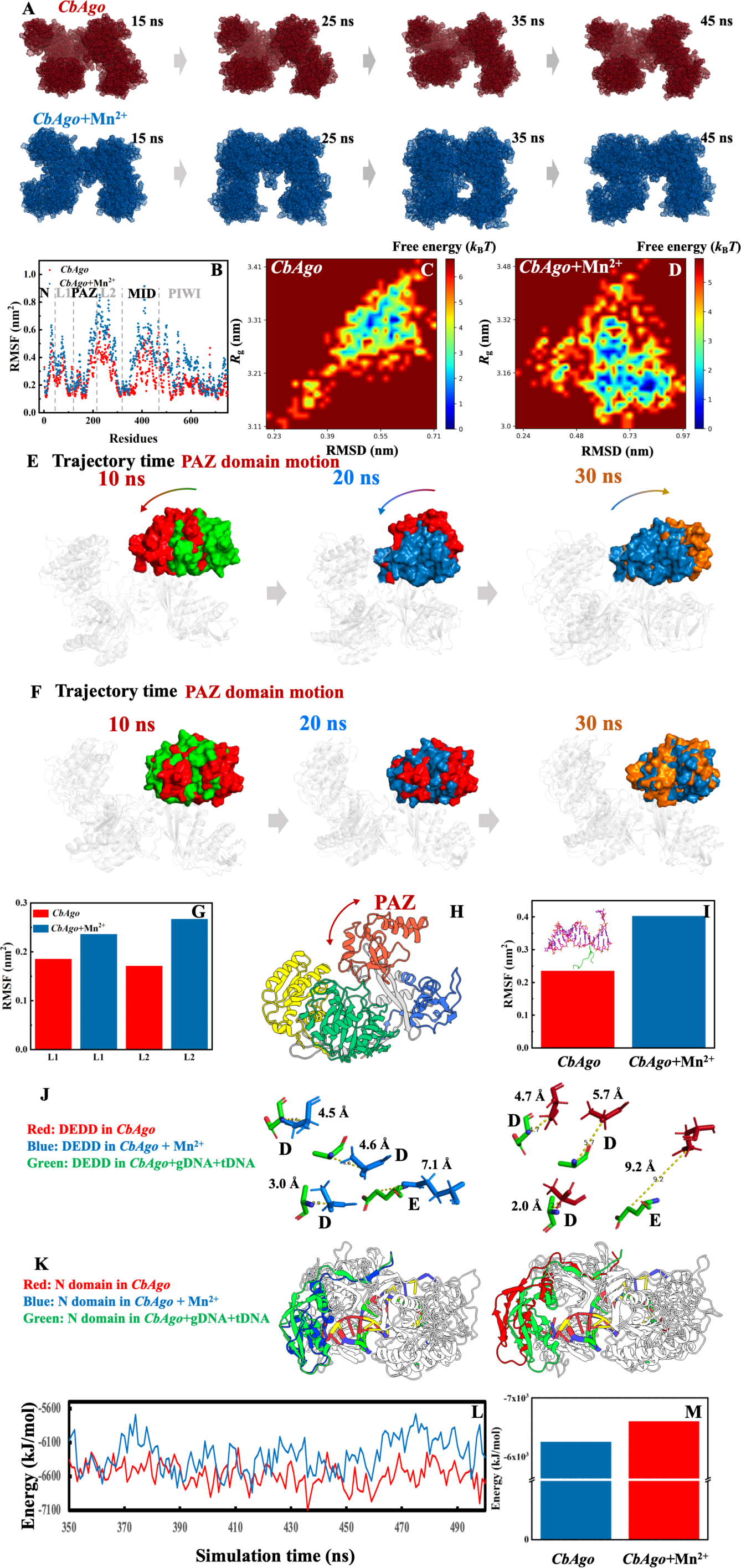
The effect of Mn^2+^ on the structural dynamics of pAgo protein derived from MD simulations. (A) The MD-derived structure of *Cb*Ago incubated with (blue) and without (red) the presence of Mn^2+^ at different simulation times. (B) Comparison of RMSF of the overall structure. Free energy landscape of *Cb*Ago incubated (C) without and (D) with Mn^2+^. RMSD is the root mean squared deviation of the structure in the first MD snapshot after equilibration. Conformational change of PAZ domain in *Cb*Ago incubated (E) with and (F) without Mn^2+^ obtained from MD simulation. Comparison of RMSF of (G) linkers and (I) catalytic loop of *Cb*Ago incubated with and without Mn^2+^. (H) Scheme of the motion of PAZ domain in pAgo protein. (J) Structural alignment of catalytic sites of *Cb*Ago incubated without (left panel) and with (right panel) Mn^2+^ with respect to that of the experimental functional conformation (PDB ID: 6qzk). Catalytic sites, DEDD, in *Cb*Ago incubated without Mn^2+^, with Mn^2+^, and in the experimental functional form are highlighted in red, blue, and green, respectively. (K) Structural alignment of N domain of *Cb*Ago incubated without (left panel) and with (right panel) Mn^2+^ with respect to that of the experimental functional conformation (PDB ID: 6QZK). N domain in *Cb*Ago incubated without Mn^2+^, with Mn^2+^, and in the experimental functional form is highlighted in red, blue, and green, respectively. Here, the protein structure is aligned by PyMOL. The alignment on *C*_alpha_ is achieved by using PyMOL software. (L) The binding energy of *Cb*Ago to gDNA/tDNA duplex over the entire simulation time derived from MD simulations. (M) The averaged binding energy of *Cb*Ago to gDNA/tDNA duplex. The initial structure of MD simulation is the protein-gDNA-tDNA tertiary complex (PDB ID: 6QZK).

#### Guide binding

Conformational mobility of the PAZ domain, upon incubation with Mn^2+^, is observed to be significantly greater than that of protein incubated without Mn^2+^ over the course of the trajectory time (see Figures 5E and F). This phenomenon can be attributed to the increased flexibility present in the Linker1 and Linker2 regions between the N domain and the MID domain, respectively (as shown in Figure 5G). The augmented conformational mobility of the PAZ domain, as indicated by the arrows in Figure 5H, could facilitate the capability of various pAgo proteins to engage with diverse populations of short DNAs of varying lengths (21, 22, 25).

#### Propagation of duplexes and cleavage of target

Figure 5I illustrates the RMSF of the glutamic loop, which houses the catalytic glutamate residue. Our analysis reveals that the glutamic loop experiences a significant increase in flexibility. The binding of the complementary strand target leads to structural modifications of pAgo proteins, including rotations of the PAZ domain and alterations in the conformations of various loops within the PIWI domain (22, 27, 34, 48). These changes propagate the nucleic acid duplex within the catalytic cleft of the pAgo protein and activate catalysis. Specifically, the glutamic loop, as shown in Figure 5I, is able to adopt distinct conformations in the presence of the target strand. It subsequently inserts into the catalytic sites and binds catalytic metal ions during the formation of the extended guide-target duplex. Therefore, the augmented conformational dynamics in the PAZ domain and catalytic loop promote the closure of the binding cleft and activate the cleavage of the target (21, 23). Furthermore, we also observed that the catalytic tetrads adjust their alignments to functional ones, thereby reducing the energy barrier for conformational change (see Figure 5J). In this context, the functional conformation of the catalytic region refers to that in the crystalline structure of the *Cb*Ago-guide-target complex (PDB ID: 4N47).

#### Target release and preparation for next round catalysis

As shown in Figure 5K, Mn^2+^ induces the position of the N domain to be adjusted closer to its functional conformation, which contributes to promoting the unwinding of the duplex of the guide and target after cleavage (28, 29). Additionally, the increased flexibility in the glutamic loop and Linkers could facilitate the unplugging of the active sites and the restoration of the PAZ domain, respectively, then regenerating the binary guide-pAgo complex for the next round of catalysis. Furthermore, we conducted an analysis of the binding energy (49) of protein to DNA in both the presence and absence of Mn^2+^. Our results indicate that the binding energy of the protein to DNA is reduced upon the addition of Mn^2+^ ions throughout the simulation time. This finding also suggests that the presence of Mn^2+^ can facilitate the release of cleaved tDNA and promote the re-adjustment of conformations (see Figures 5L and 5M).

## Discussion

Divalent cations play a fundamental role in biomacromolecule dynamics, primarily situated in ribosomes (50, 51), enzymes (30, 52, 53), and structural proteins (54). The proper function and stability of such proteins necessitate the presence of ions. Divalent ions can impart consequential effects on the conformation and functionality of proteins, which is integral to numerous biological processes like enzymatic catalysis (31), regulation of protein-protein interactions (54), and signal transduction (55). Divalent ions also play a pivotal role in the regulation of membrane transport phenomena like ion transit across the plasma membrane and intracellular movement of ions and molecules (56). In ion channels and pumps, divalent ions serve as indispensable regulators, controlling the opening and closing of these channels, thereby regulating the flux of ions into and out of cells (56). Additionally, it has been demonstrated that ions impact the activity of endonucleases. For instance, the Mg^2+^-dependent conformational rearrangements of the CRISPR-Cas12a R-loop complex are crucial for complete double-stranded DNA cleavage (52).

In the case of pAgo proteins, we unraveled the molecular mechanisms of unbound Mn^2+^ regulating the binding, cleavage, and release of products through the use of spectroscopy, biochemical assays, and MD simulation. Initially, the presence of Mn^2+^ can disrupt certain hydrogen bonds within non-secondary structures, thereby increasing the flexibility of proteins and the associated entropy, without compromising the structural integrity. Next owing to the increased flexibility of the L1 linker and L2 linker, the conformational number of the PAZ domain is significantly increased. This enhanced structural flexibility facilitates the binding of the pAgo protein to gDNA, and helps the PAZ domain recognize gDNA of varying lengths. Furthermore, the enhanced structural variability of the glutamic loop and PAZ domain aids in the propagation of the DNA duplex, preparing it for tDNA cleavage. In addition, once the holo-form arrangement is achieved by the catalytic sites and the N domain, the catalytic center in the PIWI domain can be exposed to the tDNA. This exposure allows for the cleavage of the tDNA, facilitating the release of the cleaved product. Our simulation data indicates that the energy barrier for releasing the target in this step is also lowered by the presence of Mn^2+^. The analysis of the conformational dynamics of this catalytic cycle reveals that the presence of Mn^2+^ reduces the energy barrier for conformational changes between different functional conformations. Furthermore, this result unveiled the unexpected dynamical role of unbound Mn^2+^ in increasing the conformation states of pAgo proteins, which is crucial for their catalytic functions.

Previous structural research, based on crystallography, has demonstrated that pAgo proteins must undergo significant conformational changes to implement their functions, including the rearrangement of catalytic tetrads, propagation of duplexes through shifting loops in the PIWI domain, and motion of the PAZ domain, and release of cleaved targets by adjusting the PAZ and N domains (21). Therefore, structural flexibility is crucial for the conformational arrangement needed to achieve the functionality of pAgo proteins at different stages. In our study, we found that pAgo proteins from mesophiles, thermophiles, and hyperthermophiles exhibit a significant increase in flexibility when unbounded Mn^2+^ is present.

Moreover, it has been demonstrated that pAgo proteins widely participate in hosts to defend against the invasion of nucleic acids (2). Living organisms contain DNA or RNA of different lengths and sequences, and invading nucleic acids also vary in their lengths and sequences (47). We hypothesize that pAgo proteins might need to modify their conformations to bind with different types of DNA or RNA in order to identify the invading nucleic acid. In this scenario, the flexibility of pAgo proteins, induced by Mn^2+^, may play a crucial role as it allows the protein to efficiently recognize the invader and protect the host.

The precision of nucleic acid detection methodologies is critically reliant on the accurate cleavage by pAgos. The high fidelity of such assays is underpinned by the exactness of the initial DNA cleavage process. Any deviations or inaccuracies within this fundamental step can substantially diminish the efficiency of subsequent amplification and signal readout stages (6). Our findings shed light on the role of Mn^2+^-induced flexibility in augmenting the proficiency of the pAgo protein in handling mismatches between gDNA and tDNA. We observed a correlation between increased concentrations of Mn^2+^ and an enhanced capacity of the pAgo protein to accommodate gDNA and tDNA mismatches, facilitating tDNA cleavage. The elevated presence of Mn^2+^ could enhance the protein’s conformational flexibility and various functional states, which is verified by the MD simulations. This increased structural flexibility enables the protein to adapt its conformational state to cater to a variety of mismatch scenarios, thus improving the efficiency of tDNA cleavage. These findings carry significant implications for the effectiveness of various recently developed nucleic acid detection technologies (5, 6).

## Conclusion

In the present study, we investigated how the unbound Mn^2+^-affect the dynamics, structure, and endonuclease activity of prokaryotic pAgo proteins obtained from mesophilic (*Cb*Ago, *Bl*Ago, *Pli*Ago, and *Pb*Ago), thermophilic (*Tt*Ago), and hyperthermophilic (*Pf*Ago, *Fp*Ago, and *Mf*Ago) organisms using a combination of experimental and computational methods. Our results indicate that the presence of these unbound free Mn^2+^ significantly reduces the hydrogen bonds in different pAgo proteins to release a great degree of structural flexibility while maintaining the overall structural integrity. Biochemical assays revealed that this Mn^2+^-induced structural flexibility in pAgo proteins significantly increases the substrate affinity and facilitates the binding of guide and target strands compared to the proteins without Mn^2+^. Complementary computer simulations confirmed these experimental findings and revealed that the increased flexibility of Linker1 and Linker2 promotes the conformational mobility of the PAZ domain, allowing for the recognition of guide and target strands of varying lengths. Additionally, changes in the conformations of the PAZ domain and glutamic finger promote the propagation of the duplex. By comparing the structures of pAgo proteins incubated with Mn^2+^ to those of the protein-guide-target ternary complex, we observed that the conformation of catalytic sites and N domain shifts towards a functional state, which reduces the energy cost of conformational changes required for the cleavage of target strand and the release of the products.

Our findings reveal that unbound Mn^2+^ play an important role in the endonuclease functions of pAgo proteins, and this mechanism is a general strategy employed by pAgo proteins from mesophilic, thermophilic, and hyperthermophilic organisms to enhance their catalytic functions. Unlike previous structural studies, which highlight the importance of the bound Mn^2+^ in directly participating in guide binding and target cleavage, our work discovers a role of unbound Mn^2+^ in an indirect way. We found that unbound free Mn^2+^ plays a crucial part in promoting the structural flexibility of pAgo proteins. This benefit extends not only to the binding of the protein to guide and target DNA, but also to the cleavage and release processes.

More importantly, by harnessing the influence of protein flexibility induced by unbound Mn^2+^ on the degree of mismatch between guide and target nucleic acids, we can finely tune the precision of target recognition by pAgo proteins through the modulation of Mn^2+^ concentration. This crucial characteristic expands the scope of molecular diagnostic applications, further highlighting the significance of structural flexibility in optimizing the performance of pAgo proteins.

## MATERIALS AND METHODS

### Protein expression and purification

A codon-optimized version of the *Pf*Ago, *Tt*Ago*, Fp*Ago*, Mf*Ago*, Bl*Ago*, Pb*Ago*, Cb*Ago, *Cb*Ago*M*, and *Pli*Ago gene was synthesized by Sangon Biotech (Shanghai, China), and was cloned into the pET28a plasmid to construct pEX-Ago with an N terminal His-tag. The expression plasmid was transformed into *Escherichia coli* BL21(DE3) cells. A 10 ml seed culture was grown at 37 °C in LB medium with 50 µg/ml kanamycin and was subsequently transferred to 1 L of LB in a shaker flask containing 50 µg/ml kanamycin. The cultures were incubated at 37 °C until the OD_600_ reached 0.6-0.8, and protein expression was then induced by the addition of isopropyl-D-thiogalactopyranoside (IPTG) to a final concentration of 0.5 mM, followed by incubation for 20-24 h at 18 °C. Cells were harvested by centrifugation for 30 min at 6,000 rpm, and the cell pellets were collected for later purification. The cell pellets were resuspended in lysis buffer (15 mM Tris-HCl, 500 mM NaCl, pH 7.4) and then disrupted using a High-Pressure Homogenizer at 700-800 bar for 5 min (Gefran, Italy). The lysates were centrifuged for 30 min at 12,000 rpm at 4 °C, after which the supernatants were subjected to Ni-NTA affinity purification with elution buffer (15 mM Tris-HCl, 500 mM NaCl, 250 mM imidazole, pH 7.4). Further gel filtration purification using a Superdex 200 (GE Tech, USA) was carried out with an elution buffer. The fractions resulting from gel filtration were analyzed by SDS-PAGE, and fractions containing the protein were flash frozen at −80 °C in buffer (15 mM Tris–HCl pH 7.4, 200 mM NaCl, 10% glycerin).

### Differential scanning fluorimetry

Each pAgo protein sample containing 1 μM of protein in a buffer containing 15 mM Tris-HCl (pH = 7.4) and 200 mM NaCl was prepared in triplicate and added to PCR tubes. To examine the effect of Mn^2+^ on pAgo proteins, pAgo proteins were incubated in Tris-HCl buffer (pH = 7.4) with 5 mM Mn^2+^ or 5 mM EDTA for 20 min. SYPRO Orange dye available as 5000× stock (Sigma-Aldrich) was added just before the measurement of the pAgo proteins in an appropriate amount to achieve a final concentration of the dye of 5×. The thermal denaturation of the pAgo proteins was monitored by exciting the SYPRO Orange dye at 470 nm and monitoring its fluorescence emission at 570 nm using Q-PCR (Analytikjena, Germany). The baseline correction is used by the Opticon Monitor software available on the PCR instrument.

### Nano differential scanning calorimetry

Nano differential scanning calorimetry (nanoDSC) measurements were performed by using Nano DSC instruments (TA, USA). The concentration of pAgo proteins was 0.1 mg/mL in a buffer containing 15 mM Tris-HCl (pH 7.4) and 200 mM NaCl. To examine the effect of Mn^2+^ on pAgo proteins, p*Ago* proteins were incubated in 15 mM Tris-HCl buffer (pH = 7.4) with 5 mM Mn^2+^ or 5 mM EDTA for 20 min. All the experiments were carried out at temperatures ranging from 10 to 110 °C with a heating rate of 1 °C/min and under a pressure of 3 atm. The melting curves of pAgo proteins were subtracted from the buffer scans.

### Circular dichroism spectroscopy

The CD measurements on the evolution of the secondary structure of pAgo proteins were performed in a Jasco J-1500 spectropolarimeter with a 1 mm pathlength cell. The signals in the far-UV CD region (222 nm) were monitored as a function of temperature to determine the thermal unfolding of pAgo proteins. The concentrations of pAgo proteins were 0.1 mg/mL in 1× PBS buffer (pH = 7.4) for the various measurements. The pAgo proteins were heated in the range of 15 °C to 100 °C with a 1 °C/min heating rate and equilibrated for 3 min at each temperature, and the CD data were collected at 0.5 °C intervals. For CD spectra, we used PBS buffer as the solvent instead of Tris-HCl buffer because the Tris-HCl buffer has a strong CD background, which will significantly affect the analysis of the CD signal of proteins (24).

### Small-angle X-ray scattering

Small-angle X-ray scattering (SAXS) measurement was employed to monitor the structural evolution of pAgo proteins incubated with and without Mn^2+^. In-situ synchrotron SAXS measurements were conducted at BL19U2 beamline in Shanghai Synchrotron Radiation Facility (SSRF). The X-ray wavelength was 0.103 nm. Protein samples were dissolved in a buffer containing 15 mM Tris-HCl (pH 7.4), 200 mM NaCl and 5 mM MnCl_2_ or EDTA. The concentration of samples is 0.5 mg/ml. Protein solutions were loaded into the silica cell and then gently refreshed with a syringe pump to prevent X-ray damage. The measurements were carried out at 25 °C. In order to calculate the absolute intensity of protein, the empty cell and buffer were also measured at corresponding temperatures. Two-dimensional (2D) diffraction patterns were collected by the Pilatus 2 M detector with a resolution of 1043 **×** 981 pixels of 172 μm **×** 172 μm. Twenty sequential 2D images were collected with 0.5 s exposure time per frame. The 2D scattering patterns were then integrated into one-dimensional (1D) intensity curves by using Fit2D software from European Synchrotron Radiation Facility (ESRF). Frames with no radiation damage were used for further processing. The SAXS data were compared to the atomic model by using CRYSOL software (57). *ab initio* reconstruction of protein structure by a chain-like ensemble of dummy residues is generated by GASBOR (58). P1(no symmetry) and monodisperse data mode were used in generating the GASBOR model. At least 10 iterations of GASBOR programs were independently performed to validate the models. A presentative of the most typical model obtained was used in the figures. The one-dimensional data is processed by using *ScÅtter* and ATSAS software (59). The details of SAXS data collection and analysis are summarized in Table S3.

### Molecular dynamics simulations

The initial structures of *Cb*Ago and *Tt*Ago for simulations were taken from PDB crystal structures 6QZK and 4N47, respectively. The initial structures of *Bl*Ago and *Fp*Ago for simulations were taken from AlphaFold2 (60). The initial structure of *Cb*Ago*-*gDNA-tDNA for simulations was taken from PDB crystal structures 6QZK. Protein and a large number of water molecules were filled in a cubic box (see Figure S5). For proteins and complexes incubated with the Mn^2+^ system, 6 Mn^2+^ were added into the simulation box to mimic the experimental concentration, and 32 chlorine counter ions were added to keep the system neutral in charge. For proteins and complexes in an aqueous solution, 16 chlorine counter ions were added to keep the system neutral in charge. The CHARMM36m force field (61) was used for proteins, complex, and Mn^2+^, and the CHARMM-modified TIP3P model was chosen for water. The simulations were carried out at 298 K. After the 4000-step energy-minimization procedure, the systems were heated and equilibrated for 100 ps in the NVT ensemble and 500 ps in the NPT ensemble. The 100-ns production simulationswere carried out at 1 atm with the proper periodic boundary condition, and the integration step was set to 2 fs. Fig. S3 shows the 100 ns profile of potential energy as a function of MD trajectory time for *Cb*Ago incubated with and without Mn^2+^. It was clear that the equilibration procedure was sufficient for minimizing the energy of protein structures. The covalent bonds with hydrogen atoms were constrained by the LINCS algorithm (62). Lennard-Jones interactions were truncated at 12 Å with a force-switching function from 10 to 12 Å. The electrostatic interactions were calculated using the particle mesh Ewald method (63) with a cutoff of 12 Å on an approximately 1 Å grid with a fourth-order spline. The temperature and pressure of thesystem are controlled by the velocity rescaling thermostat (64) and the Parrinello-Rahman algorithm (65), respectively. All MD simulations were performed using GROMACS 2020.4 software packages. Representative simulation snapshots of the systems are given in Figure S5.

### Single-strand DNA cleavage assay

For standard activity assays, cleavage experiments were performed in a 5:1:1 molar ratio (protein:guide:target). First, 5 µM pAgo protein was mixed with a synthetic 1 μM gDNAguide in the reaction buffer (15 mM Tris-HCl (pH 7.4), 200 mM NaCl, different concentrations of MnCl_2_ or 5 mM EDTA). The solution was then pre-incubated at 37 ℃ for 20 min. After pre-incubation, 1 μM tDNA, which was labeled with the fluorescent group 6-FAM at the 5’-end and the quencher BHQ1 at the 3’-end, was added to the mixture. The cleavage experiments were performed at 37 °C. All experiments were performed in triplicate, and the fluorescence signals were traced by the quantitative real-time PCR QuantStudio 5 (Thermo Fisher Scientific, USA) with λ_ex_ = 495 nm and λ_em_ = 520 nm. The results were analyzed by QuantStudio^TM^ Design & Analysis Software v1.5.1. All the gDNA and tDNA used for cleavage are listed in Table S4.

### Fluorescence Polarization Assay

To determine the apparent dissociation constant (*K*_d_) for proteins binding to either gDNA or tDNA, a fluorescence polarization assay was conducted using a multifunctional enzyme-linked immunosorbent assay plate reader (Spark, Tecan). A solution was prepared by combining 5 nM of 3’ 6-FAM labeled guide or target DNA with proteins across a concentration range of 0 to 1500 nM in a reaction buffer (15 mM Tris-HCl at pH 7.4, 200 mM NaCl, and 5 mM MnCl_2_). This mixture was incubated at 37°C for 1 hour and subsequently transferred to a light-protected 96-well ELISA plate. The degree of polarization was measured using the Spark Tecan plate reader, employing an excitation wavelength of 485 nm and an emission wavelength of 525 nm. All experiments were independently conducted three times. The binding percentages were analyzed using Microsoft Excel and Prism 8 (GraphPad) software. The data was fitted with the Hill equation, incorporating a Hill coefficient of 2 to 2.5.

## Data availability

All the data are presented in SI.

## Acknowledgments

We would like to thank the Instrumental Analysis Center of Shanghai Jiao Tong University for assistance with thermal performance tests via nanoDSC and CD. We would like to thank Dr. Na Li from BL19U2 beamline of Shanghai Synchrotron Radiation Facility (SSRF) for the help with synchrotron X-ray measurements on No. h20pr0008.

