## Supporting information for "Mn^2+^-Induced Structural Flexibility Enhances the Entire Catalytic Cycle and the Cleavage of Mismatches in Prokaryotic Argonaute Proteins"

**This PDF file includes:**

Figure S1-S23

Table S1-S6

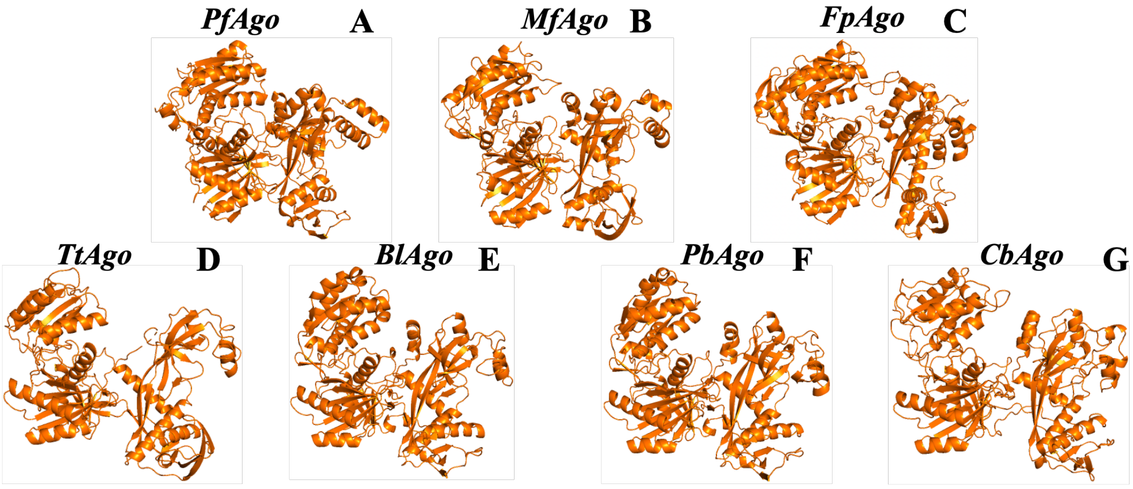

**Figure S1.** Atomic structure of (A) *Pf*Ago, (B) *Mf*Ago, (C) *Fp*Ago, (D) *Tt*Ago, (E) *Bl*Ago, (F) *Pb*Ago, and (G) *Cb*Ago. The atomic structures of *Pf*Ago, *Tt*Ago, and *Cb*Ago are obtained from the crystal structure PDB ID: 1Z25, 4N47, and 6QZK, respectively. The atomic structures of *Mf*Ago, *Fp*Ago, *Bl*Ago, *Pb*Ago are obtained from Alphafold2.

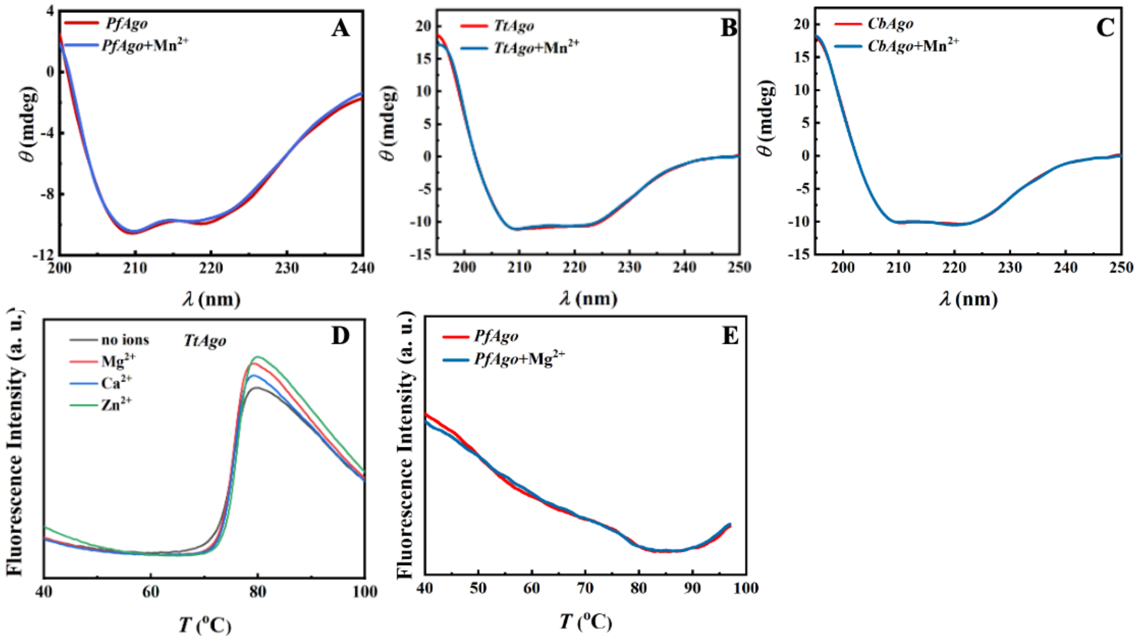

**Figure S2.** The CD spectra of (A) *Pf*Ago, (B) *Tt*Ago, and (C) *Cb*Ago incubated with and without $Mn^{2+}$ at their respective physiological temperatures. The DSF of (D) *Tt*Ago and (E) *Pf*Ago incubated with and without divalent cations.

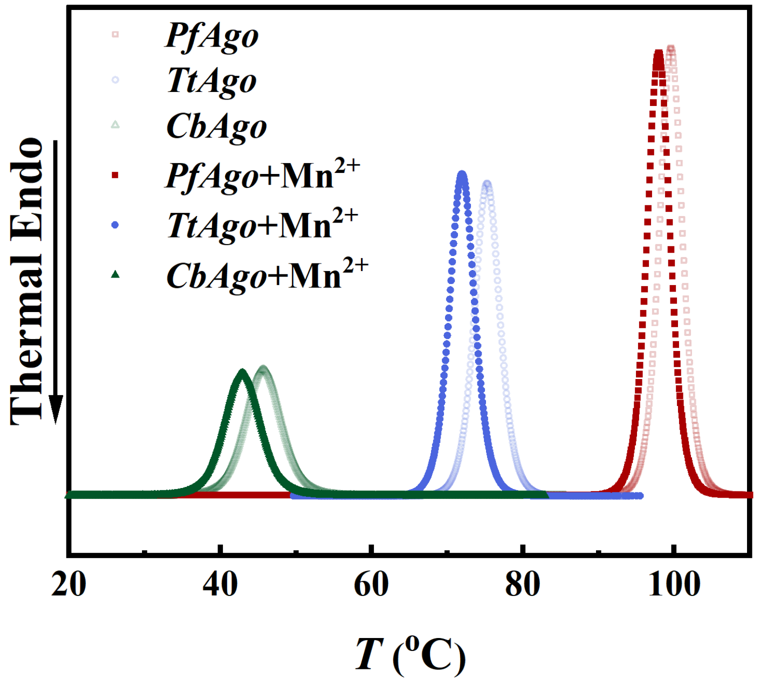

**Figure S3.** Thermal denature curves of pAgo proteins incubated with and without $Mn^{2+}$ determined by nanoDSC.

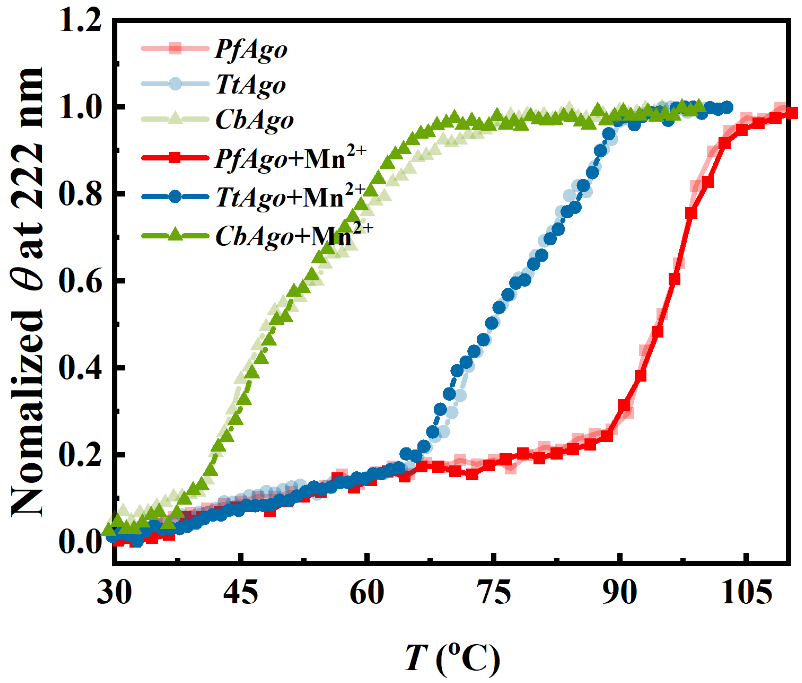

**Figure S4.** Thermal denature curves of pAgo proteins incubated with and without $Mn^{2+}$ determined by CD.

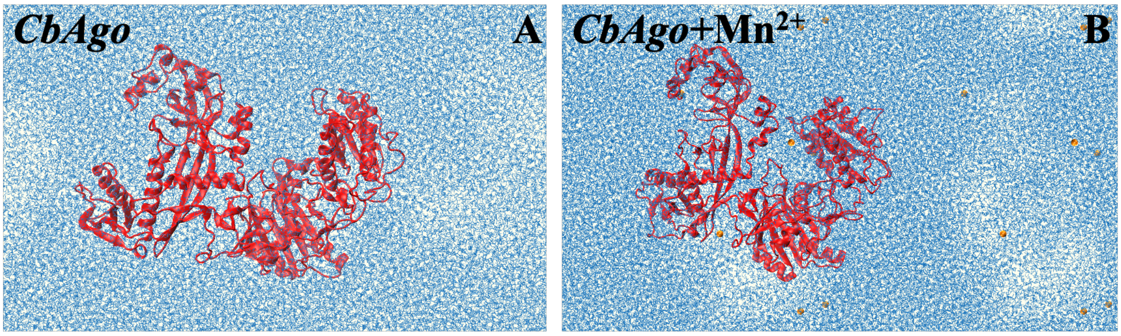

**Figure S5.** A snapshot of the MD simulation of *Cb*Ago incubated (A) without and (B) with $Mn^{2+}$ in solution. The golden spheres represent $Mn^{2+}$.

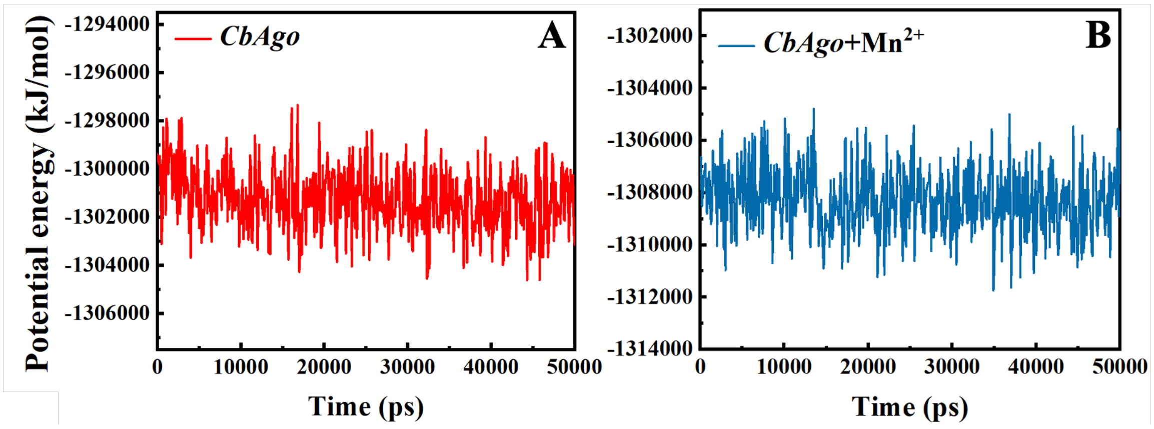

**Figure S6**. The potential energy of protein as a function of MD simulation time of *Cb*Ago incubated (A) without and (B) with $Mn^{2+}$. The simulation time is set as 50 ns.

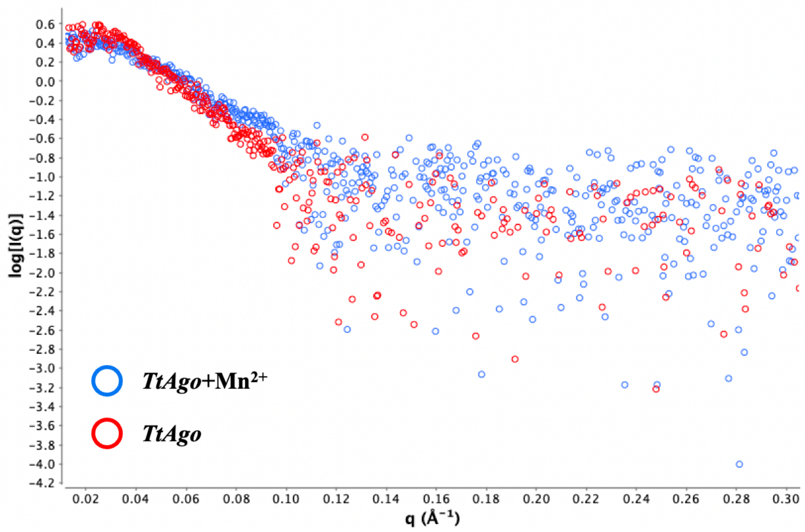

**Figure S7.** SAXS curves of *Tt*Ago incubated with and without $Mn^{2+}$.

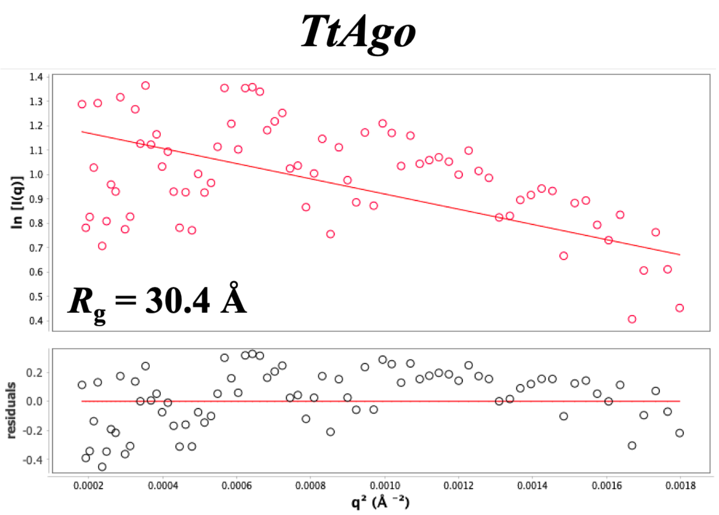

**Figure S8.** Guinier plots of *Tt*Ago. The lower insets show the error-weighted residual difference plots for the Guinier fitting.

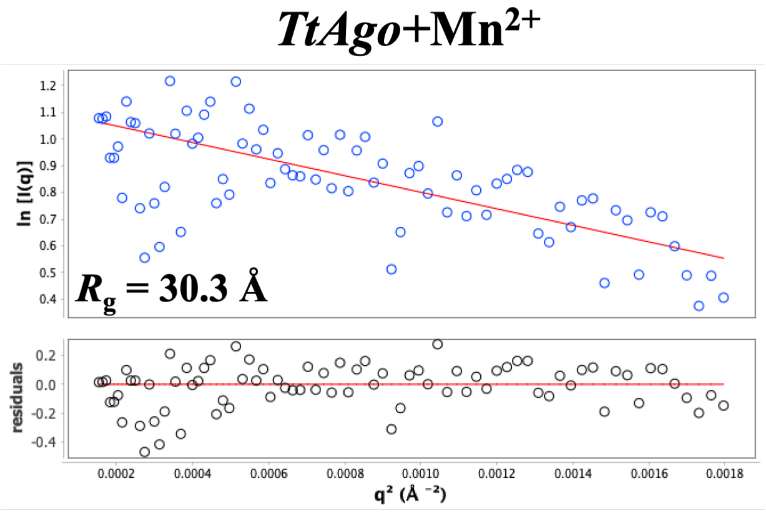

**Figure S9.** Guinier plots of *Tt*Ago incubated with $Mn^{2+}$. The lower insets show the error-weighted residual difference plots for the Guinier fitting.

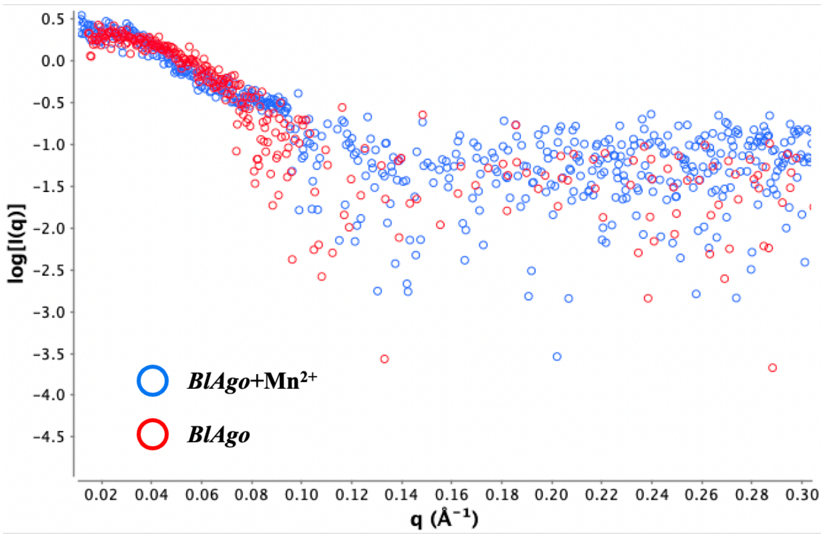

**Figure S10.** SAXS curves of *Bl*Ago incubated with and without $Mn^{2+}$.

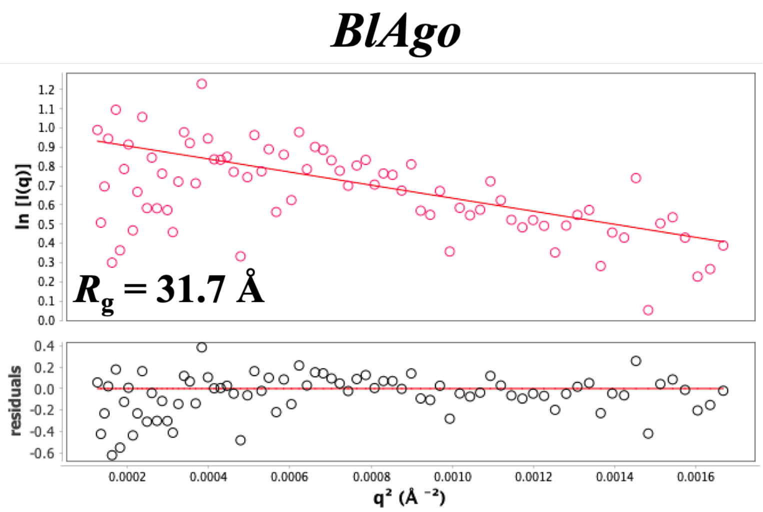

**Figure S11.** Guinier plots of *Bl*Ago. The lower insets show the error-weighted residual difference plots for the Guinier fitting.

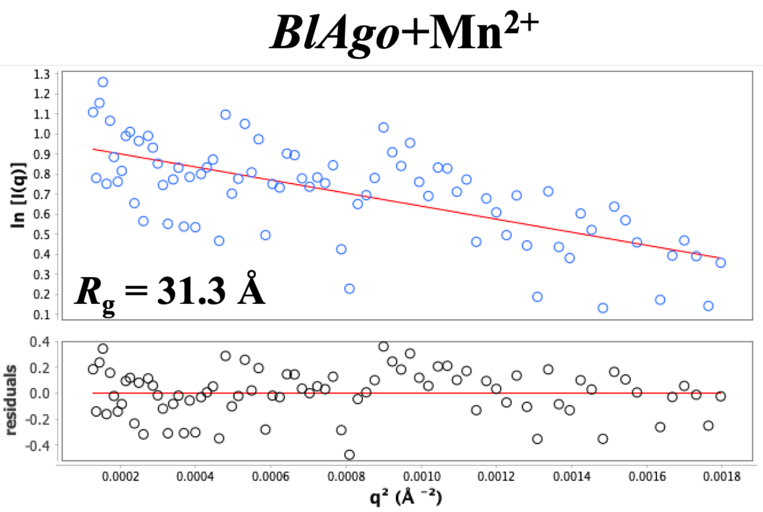

**Figure S12.** Guinier plots of *Bl*Ago incubated with $Mn^{2+}$. The lower insets show the error-weighted residual difference plots for the Guinier fitting.

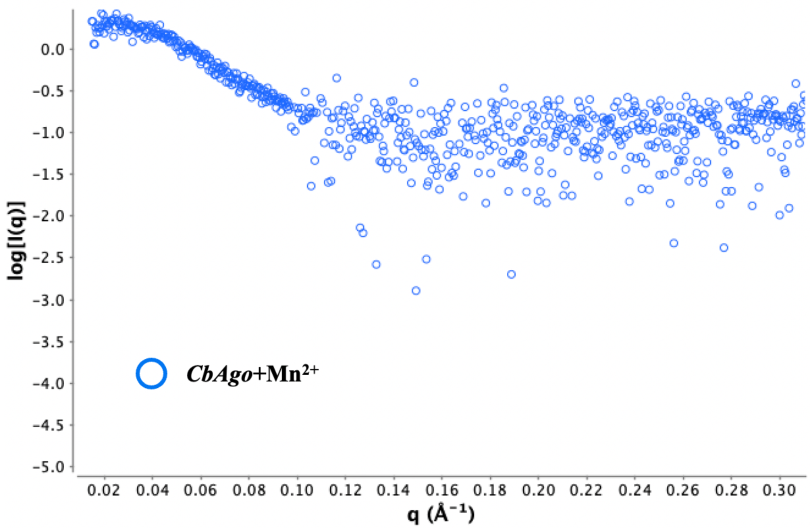

**Figure S13.** SAXS curve of *Cb*Ago incubated with $Mn^{2+}$.

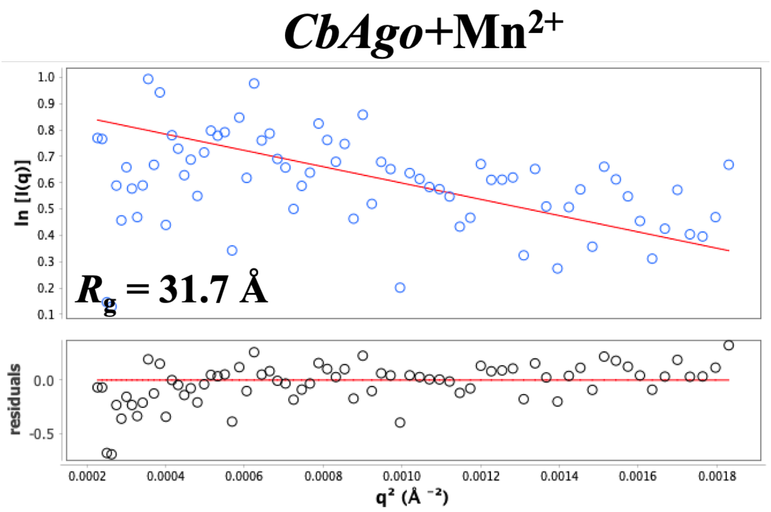

**Figure S14.** Guinier plots of *Cb*Ago incubated with $Mn^{2+}$. The lower insets show the error-weighted residual difference plots for the Guinier fitting.

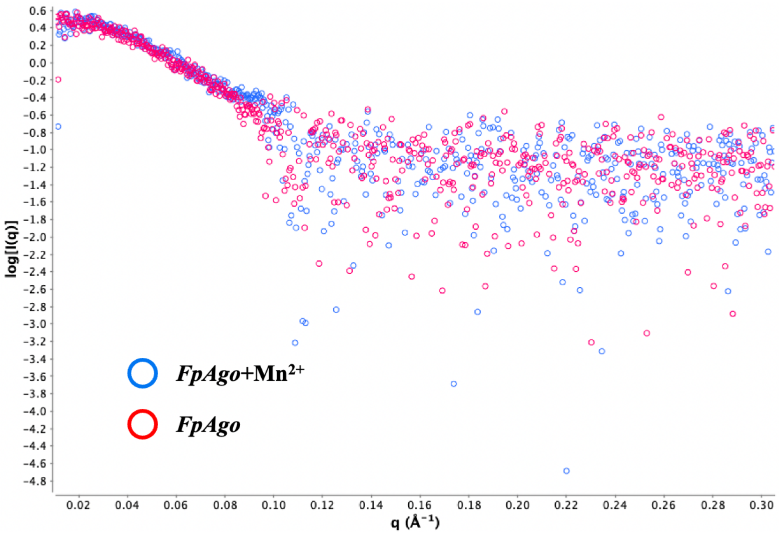

**Figure S15.** SAXS curves of *Fp*Ago incubated with and without $Mn^{2+}$.

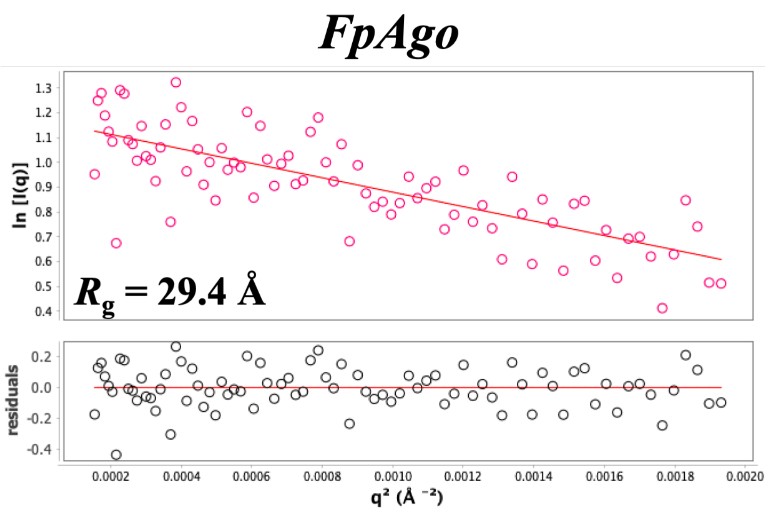

**Figure S16.** Guinier plots of *Fp*Ago. The lower insets show the error-weighted residual difference plots for the Guinier fitting.

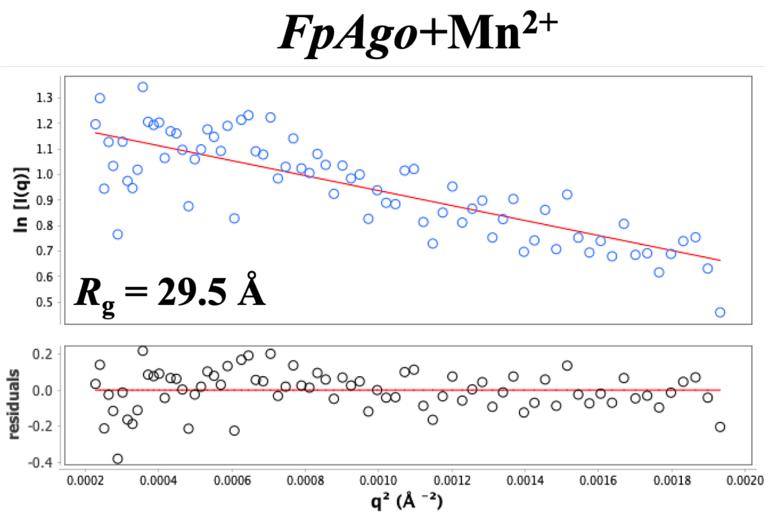

**Figure S17**. Guinier plots of *Fp*Ago incubated with $Mn^{2+}$. The lower insets show the error-weighted residual difference plots for the Guinier fitting.

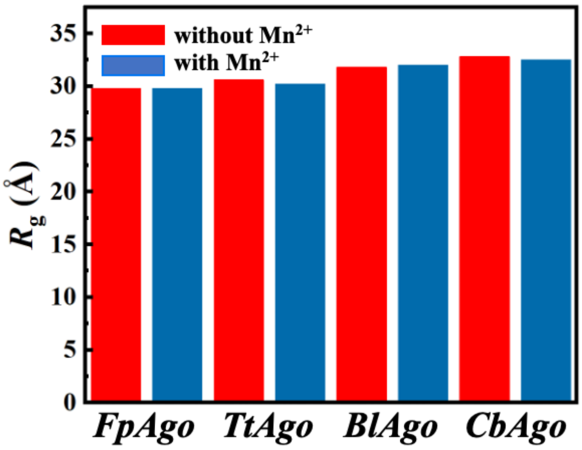

**Figure S18.** The radius of gyration ($R_g$) of *Fp*Ago, *Tt*Ago, *Bl*Ago, and *Cb*Ago incubated with and without $Mn^{2+}$ derived from MD simulations.

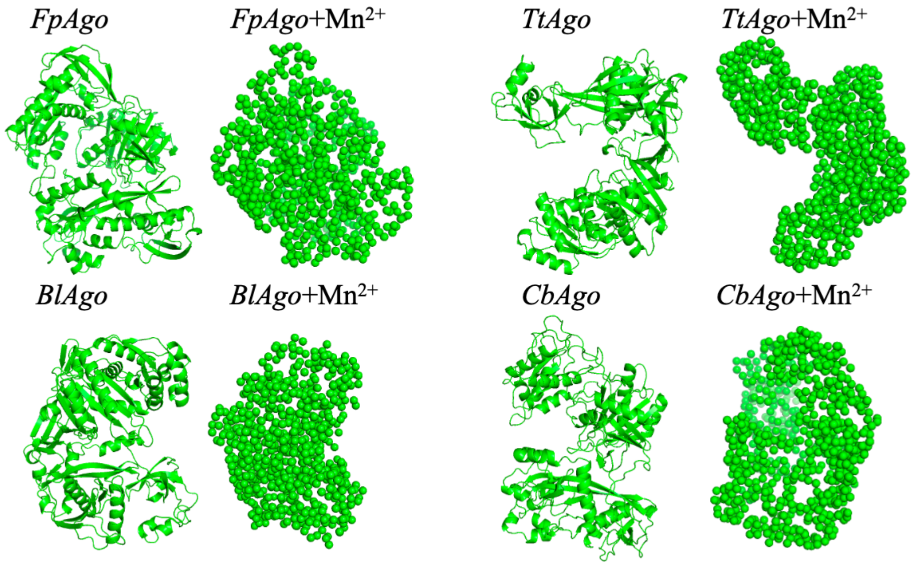

**Figure S19.** The *ab initio* structures of (A) *Fp*Ago, (B) *Tt*Ago, (C) *Bl*Ago, and (D) *Cb*Ago.

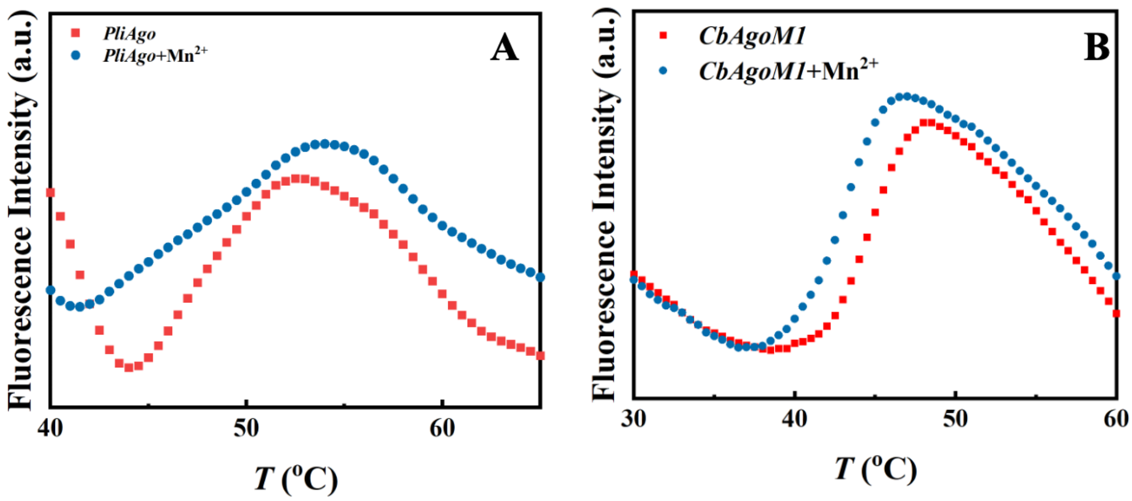

**Figure S20.** DSF measurements of (A) *Pli*Ago and (B) *Cb*Ago*M1*.

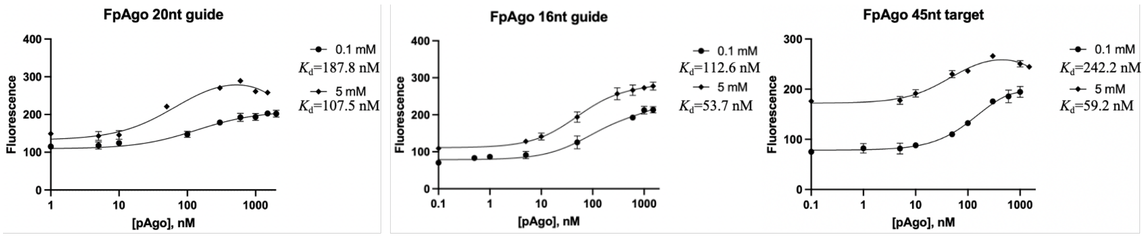

**Figure S21.** Fluorescence polarization assay of the tDNA and gDNA of varying lengths at different concentrations of $Mn^{2+}$. The results from three independent experiments were quantified. Error bars represent the standard deviations of three independent experiments.

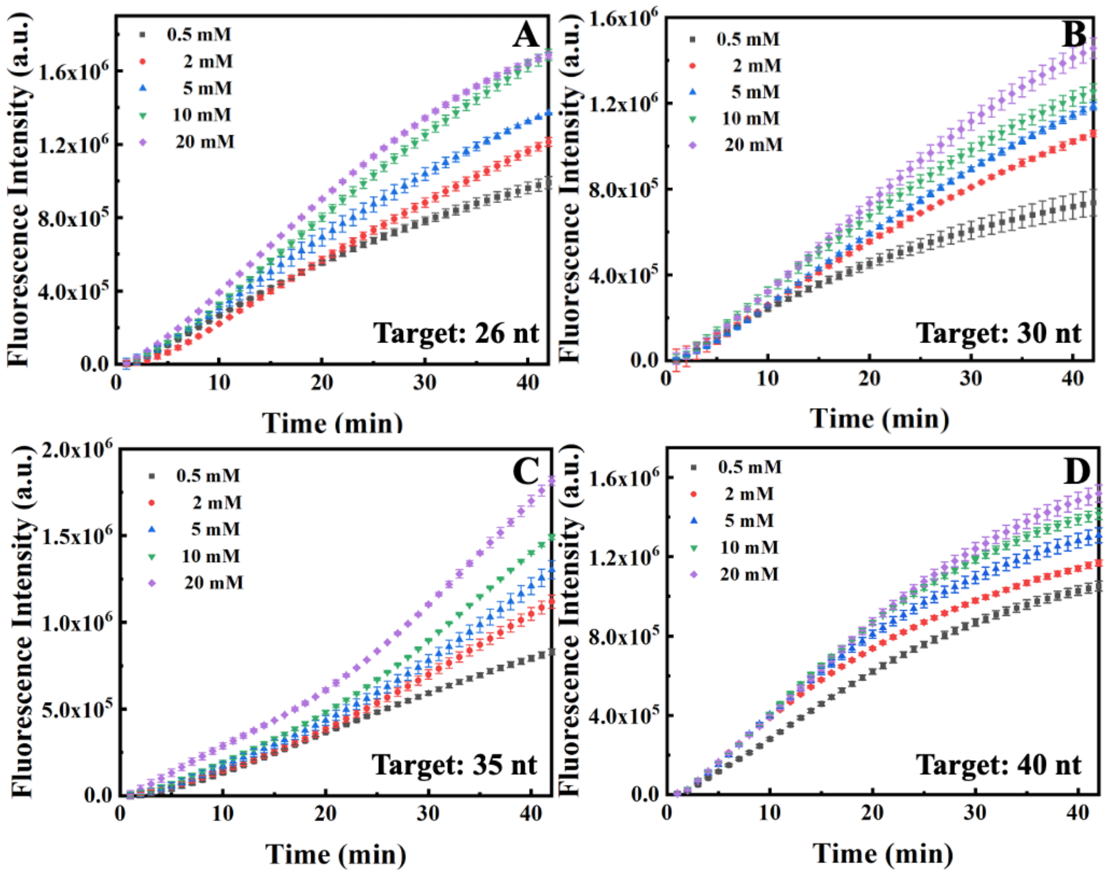

**Figure S22.** The role of the $Mn^{2+}$-dependent structural dynamics of pAgo proteins for cleavage target DNA of varying length. Cleavage assays with gDNA and tDNA of varying lengths at different concentrations of $Mn^{2+}$. *Pli*Ago preloaded with 21 nt gDNA cleaves (A) 26 nt, (B) 30 nt, (C) 35 nt, and (D) 40 nt tDNA. In all experiments, protein, guide, and target were mixed at a 5:1:1 molar ratio and incubated at 37 °C. The reaction buffer without $Mn^{2+}$ contains 10 mM EDTA. Three samples were used for each experimental condition. The results from three independent experiments were quantified. Error bars represent the standard deviations of three independent experiments. The detailed procedure is presented in the Materials and Methods. The nucleotide sequences of the gDNA and tDNA are presented in Table S4.

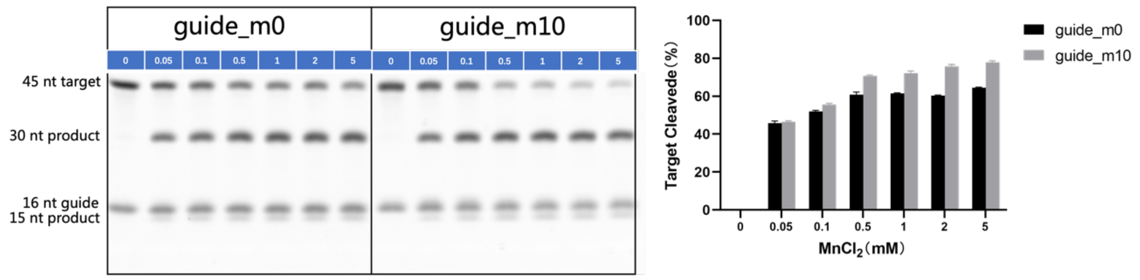

**Figure S23.** Left panel: Gel detection results for the designed mismatched gDNA-directed cleavage on tDNA at different concentrations of $Mn^{2+}$. Quantifying the effects of incubating with $Mn^{2+}$ on tDNA cleavage obtained from the left panel. *Cb*Ago is used here. The results from three independent experiments were quantified. Error bars represent the standard deviations of three independent experiments.

**Table S1.** The relative contents of different secondary structures in pAgo proteins derived from MD simulations, as defined by mass fraction.

| Protein | Alpha-helix (%) | Beta-sheet (%) | Turn and loops (%) |
|---|---|---|---|
| ***Cb*Ago+$Mn^{2+}$** | 29 | 25 | 46 |
| ***Cb*Ago** | 29 | 25 | 46 |
| ***Tt*Ago+$Mn^{2+}$** | 31 | 31 | 38 |
| ***Tt*Ago** | 31 | 31 | 38 |
| ***Fp*Ago+$Mn^{2+}$** | 33 | 28 | 39 |
| ***Fp*Ago** | 33 | 28 | 39 |
| ***Bl*Ago+$Mn^{2+}$** | 30 | 27 | 43 |
| ***Bl*Ago** | 30 | 27 | 43 |

**Table S2.** $M_w$ of *Cb*Ago, *Tt*Ago, *Bl*Ago, and *Fp*Ago estimated from SAXS. $M_W^{expected}$ is obtained from the amino acids sequence.

| Protein | $M_W^{experimental}$ (kDa) | $M_W^{expected}$ (kDa) |
|---|---|---|
| *Cb*Ago | 91 | 86 |
| *Cb*Ago+$Mn^{2+}$ | 88 | |
| *Tt*Ago | 78 | 77 |
| *Tt*Ago+$Mn^{2+}$ | 79 | |
| *Bl*Ago | 72 | 80 |
| *Bl*Ago+$Mn^{2+}$ | 78 | |
| *Fp*Ago | 90 | 83 |
| *Fp*Ago+$Mn^{2+}$ | 84 | |

**Table S3.** SAXS data collection and analysis.

| | *FpAgo* | *FpAgo*+ $Mn^{2+}$ | *TtAgo* | *TtAgo*+ $Mn^{2+}$ | *CbAgo* | *CbAgo* + $Mn^{2+}$ | *BlAgo* | *BlAgo*+ $Mn^{2+}$ |
|---|---|---|---|---|---|---|---|---|
| **Data acquisition** | | | | | | | | |
| Beamline facility | BL19U2-SSRF | | BL19U2-SSRF | | BL19U2-SSRF | | BL19U2-SSRF | |
| Wavelength (nm) | 0.103 | | 0.103 | | 0.103 | | 0.103 | |
| Sample-to-detector distance (m) | 3 | | 3 | | 3 | | 3 | |
| Detector | Pilatus 2M | | Pilatus 2M | | Pilatus 2M | | Pilatus 2M | |
| **Overall parameter** | | | | | | | | |
| $R_g$ (Å) (from P(r)) | 2.95 | 2.95 | 3.0 | 3.1 | 3.20 | 3.20 | 3.14 | 3.17 |
| $R_g$ (Å) (from Guinier) | 2.94 | 2.95 | 3.04 | 3.03 | 3.20 | 3.17 | 3.17 | 3.13 |
| Porod volume estimate ($nm^3$) | 148 | 150 | 122 | 121 | 185 | 188 | 195 | 192 |
| Oligomeric state | monomer | | monomer | | monomer | | monomer | |
| **Software** | | | | | | | | |
| SAXS data integration | *ScÅtter* | | *ScÅtter* | | *ScÅtter* | | *ScÅtter* | |
| Ab initio modeling | GASBOR | | GASBOR | | GASBOR | | GASBOR | |
| Simulated SAXS | CRYSOL | | CRYSOL | | CRYSOL | | CRYSOL | |

**Table S4.** List of nucleic acids sequences for binding and cleavage assay used in this study

| Oligonucleotide name | Sequence (5'-3') | Description |
|---|---|---|
| gDNA 21 nt | 5'P-TGAGGTAGTAGGTTGTATAGT | |
| tDNA 26 nt | 5'FAM-ATATACTATACAACCTACTACCTCGT-3'BHQ1 | 5'FAM labeled tDNA, 3'BHQ1 |
| tDNA 30 nt | 5'FAM-ATATACTATACAACCTACTACCTCGTATAA-3'BHQ1 | 5'FAM labeled tDNA, 3'BHQ1 |
| tDNA 35 nt | 5'FAM-ATATACTATACAACCTACTACCTCGTATAAATTTT-3'BHQ1 | 5'FAM labeled tDNA, 3'BHQ1 |
| tDNA 40 nt | 5'FAM-ATATACTATACAACCTACTACCTCGTATAAATTTTAAAT-3'BHQ1 | 5'FAM labeled tDNA, 3'BHQ1 |
| tDNA 45 nt | 5'FAM-ATATACTATACAACCTACTACCTCGTATAAATTTTAAATAAAT-3'BHQ1 | 5'FAM labeled tDNA, 3'BHQ1 |

**Table S5.** List of amino acids sequences of pAgo proteins used in this study

| p*Ago* proteins | Sequence | Description |
|---|---|---|
| *Cb*Ago | MANNLTFEAFEGIGQLNELNFYKYRLIGKGQIDNVHQAIWSVK<br>YKLQANNFFKPVFVKGEILYSLDELKVIPEFENVEVILDGNIILSI<br>SENTDIYKDVIVFYINNALKNIKDITNYRKYITKNTDEIICKSILTT<br>NLKYQYMKSEKGFKLQRKFKISPVVFRNGKVILYLNCSSDFSTD<br>KSIYEMLNDGLGVVGLQVKNKWTNANGNIFIEKVLDNTISDPG<br>TSGKLGQSLIDYYINGNQKYRVEKFTDEDKNAKVIQAKIKNKT<br>YNYIPQALTPVITREYLSHTDKKFSKQIENVIKMDMNYRYQTLK<br>SFVEDIGVIKELNNLHFKNQYYTNFDFMGFESGVLEEPVLMGA<br>NGKIKDKKQIFINGFFKNPKENVKFGVLYPEGCMENAQSIARSIL<br>DFATAGKYNKQENKYISKNLMNIGFKPSECIFESYKLGDITEYKA<br>TARKLKEHEKVGFVIAVIPDMNELEVENPYNPFKKVWAKLNIPS<br>QMITLKTTEKFKNIVDKSGLYYLHNIALNILGKIGGIPWIIKDMP<br>GNIDCFIGLAVGTREKGIHFPACSVLFDKYGKLINYYKPTIPQSG<br>EKIAETILQEIFDNVLISYKEENGEYPKNIVIHRAGFSRENIDWYK<br>EYFDKKGIKFNIIEVKKNIPVKIAKVVGSNICNPIKGSYVLKNDK<br>AFIVTTDIKDGVASPNPLKIEKTYGDVEMKSILEQIYSLSQIHVGS<br>TKSLRLPITTGYADKICKAIEYIPQGVVDNRLFFL | Wild type |
| *Cb*Ago*M* | MNNLTFEAFEGIGQLNELNFYKYRLIGKGQIDNVHQAIWSVKY<br>KLQANNFFKPVFVKGEILYSLDELKVIPEFENVEVILDGNIILSIS<br>ENTDIYKDVIVFYINNALKNIKDITNYRKYITKNTDEIICKSILTT<br>NLKYQYMKSEKGFKLQRKFKISPVVFRNGKVILYLNCSSDFSTD<br>KSIYEMLNDGLGVVGLQVKNKWTNANGNIFIEKVLDNTISDPG<br>TSGKLGQSLIDYYINGNQKYRVEKFTDEDKNAKVIQAKIKNKT<br>YNYIPQALTPVITREYLSHTDKKFSKQIENVIKMDMNYRYQTLK<br>SFVEDIGVIKELNNLHFKNQYYTNFDFMGFESGVLEEPVLMGA<br>NGKIKDKKQIFINGFFKNPKENVKFGVLYPEGCMENAQSIARSIL<br>DFATAGKYNKQENKYISKNLMNIGFKPSECIFESYKLGDITEYKA<br>TARKLKEHEKVGFVIAVIPRMNELEVENPANPFCKVWAKLNIPS<br>QMIHLKSTEKFKNIVDKSGLYYLHNIALNILGGIGGIPWIIKDMP<br>GNIDCFIGLDVGTREKGIHFPACSVLFDKYGKLINYYKPTIPQSG<br>EKIAETILQEIFDNVLISYKEENGEYPKNIVIHRDGFSRENIDWYK<br>EYFDKKGIKFNIIEVKKNIPVKIAKVVGSNICNPIKGSYVLKNDK<br>AFIVTTDIKDGVASPNPLKIEKTYGDVEMKSILEQIYSLSQIHVGS<br>TKSLRLPITTGYADKICKAIEYI | Mutant |
| *Pb*Ago | MNTPLTHYVLTEWESDTNTNVLHIHLYTLPVRNVFEQHKENGN<br>ACFDLRKLNRSLIIDFYDQYIVSWQPIENWGEYTFTQHEYRSINP<br>TILAERAILERLLLRTIESVQPKKEIAAGSRKFTWLKAEKVVENI<br>SIHRVIQCDVTVDYAGKISVGFDLNHSYRTNESVYDLMKSNAIF<br>KGDRVIDIYNNLHYEFVEISNSTINDSIPELNQSVVNYFTKERKQ<br>AWKVDKLEQSMPVVYLKAFNGSRIAYAPAMLQKELTFESLPTN<br>VVRQTSEIFKQNANQKIKTLLDEIQKILARTDKIKFNKQKLLVQ<br>QAGYEILELSNPNLQFGKNVTQTQLKYGLDKGGVVASKPLSINL<br>LVYPELIDTKLDVINDFNDKLNALSHKWGVPLSILKKSGAYRNR<br>PIDFTNPHQLAILLKELTKNLFQELTLVIIPEKISGMWYDLVKKEF<br>GGNSSVPTQFITIETLQKANDYILGNLLLGLYSKSGIQPWILNSPL<br>SSDCFIGLDVSHEAGRHSTGIVQVVGKDGRVLSSKANTSNEAGE<br>KIRHETMCQIVYSAIDQYQQHYNERPKHVTFHRDGFCREDLLS<br>LDEVMNSLDVQYDMVEIIKKTNRRMALTVGKQGWETKPGLCY<br>LKDESAYLIATNPHPRVGTAQPIKIIKKKGSLPIEAIIQDIYHLSFM<br>HIGSLLKCRLPITTYYADLSSTFFNRQWLPIDSGEALHFV | Wild type |
| *Pli*Ago | MTLETTLFPLEGLEGLTASYQLYAVKGLSGLDETEYHKNVNLLV<br>RRLSFSMKAPFVALSRDGEQFIAVPNYVTEFPVDHRVVRAMVK<br>LVPTGEPLNLRFDAADDEYDGLRLRYLDFVLQQPLFANHHLWQ<br>PGSGQPFFHKKPLKRLDDVDLYDGVSVRAAKHPEGGFGIVCDA<br>RSKFITHTPIGARADRKRLGKLINRSCLYKMGDHWYQFRIDAVS<br>DWKVGEPSLFEGNVPISLAQQLVRTAGNAAPKSIIDLDPEGGAL<br>EYFTSTNERRMAPAELCFLIEDTHGRRAAKLQRQTILSPSERRAR | Wild type |

| | | |
|---|---|---|
| | VNGFIRRYLSELNIGGAKLSAGARAHAFFTETHMPPALSFGNGT<br>VLAPDTSKDRFQAMQEYSSMRRTMMLDKKVGFFHQDVFPPQT<br>LLLPESVKKSWGPAFASDFVGTVQELYPAGGYRPEIIEYRDKAY<br>GGGVPGQMKALLEVAERGEIKSGDVLVMLHRINGAPRAQDKL<br>AAMVCNEFEKRFGKRVQVIHSDSPGRGYKRIFKNDKPTYVQQR<br>GRGVNIKGYLKGAALNKVCLGNSRWPFVLRDPLNADVTIGIDV<br>KNNMAVFTMVAEGGRIVRVQRSRSRQREQLLESQVTQVITEML<br>SKELPEIKKQVQRVVIHRDGRAWPAEIAGARKTFADMAESGLIA<br>VDADVSVFEVLKSSPAPLRLFSFEEPTQENPKGVINPVLGSWLK<br>LSENDGYICTTGAPLLLQGTADPLHVRKAFGPMAIEDALKDVF<br>DLSCLTWPKPDSCMRLPLTIKLCDIALFDDAAEYDVDVVRFAD<br>GNTGEASA | |
| *Bl*Ago | MNDFQVLTEWTLDTRATEFIIHIYTMPINDLNKSHTESYELVKTL<br>RRLNNTKEIVFYEQQIAALTKVENWGHYTRGDYQHRCIKLSVP<br>RERELLQRLLLKVVGARQPKKVYAEGGLFINASPQYKIENINIH<br>EALNLDFSVTQEGLIIVGFDFTHKLYYKDTLLEFVKRNQIQKGD<br>RVIDPIYHISYVYDEVAPYTVSESSPYLGQSILQYYKNKDWILKK<br>LNTDMSVVHVRNKENKIFPYAPVFLKKECSLSSLDQRIVTKINR<br>VIKLGPNEKMTKSIREVQNILSRFEVLRLNKNNLLVSNQGYKIK<br>YFPVPSLRFGKNVTSKSLKNGLIKGGVVDPRNLELAYFIDPVIIK<br>NNTESIEVFMKNLEEKSKQLGVPLQRLKKGRSFYNQLDTQMFS<br>SPNELVLGLKKIAKEFNCLTVVITTQQNIDKCYGAIKKEFGGNY<br>DIPTQFVTADTAKEKNDYILLNILLGIYAKASIQSWILKEPLHSDC<br>FIGLDVSHEENRHSTGLVQVVGKDGRVLSSKAMSTIESGERIRD<br>ETMKEIVYEAIHSYENQYGYRPKHVTFHRDGFCRENIDNIEYIL<br>KNLGVLFDYVEVIKKSNRRMANFLSKEEGWKTEIGASYLKDDL<br>AYLCSTAPGKQVGMAIPVKITQVTGHLTMSAIVSDIFNLSHMHV<br>GSLIKSRLPISTYYADLSSTFFNRGWISSRSNGLQFV | Wild type |
| *Tt*Ago | MNHLGKTEVFLNRFALRPLNPEELRPWRLEVVLDPPPGREEVYPLLAQVARRAGGVT<br>VRMGDGLASWSPPEVLVLEGTLARMGQTYAYRLYPKGRRPLDPKDPGERSVLSALAR<br>RLLQERLRRLEGVWVEGLAVYRREHARGPGWRVLGGAVLDLWVSDSGAFLLEVDPAY<br>RILCEMSLEAWLAQGHPLPKRVRNAYDRRTWELLRLGEEDPKELPLPGGLSLLDYHA<br>SKGRLQGREGGRVAWVADPKDPRKPIPHLTGLLVPVLTLEDLHEEEGSLALSLPWEE<br>RRRRTREIASWIGRRLGLGTPEAVRAQAYRLSIPKLMGRRAVSKPADALRVGFYRAQ<br>ETALALLRLDGAQGWPEFLRRALLRAFGASGASLRLHTLHAHPSQGLAFREALRKAK<br>EEGVQAVLVLTPPMAWEDRNRLKALLLREGLPSQILNVPLREEERHRWENALLGLLA<br>KAGLQVVALSGAYPAELAVGFDAGGRESFRFGGAACAVGGDGGHLLWTLPEAQAGER<br>IPQEVVWDLLEETLWAFRRKAGRLPSRVLLLRDGRVPQDEFALALEALAREGIAYDL<br>VSVRKSGGGRVYPVQGRLADGLYVPLEDKTFLLLTVHRDFRGTPRPLKLVHEAGDTP<br>LEALAHQIFHLTRLYPASGFAFPRLPAPLHLADRLVKEVGRLGIRHLKEVDREKLFF<br>V | Wild type |
| *Pf*Ago | MKAKVVINLVKINKKIIPDKIYVYRLFNDPEEELQKEGYSIYRLA<br>YENVGIVIDPENLIIATTKELEYEGEFIPEGEISFSELRNDYQSKLV<br>LRLLKENGIGEYELSKLLRKFRKPKTFGDYKVIPSVEMSVIKHD<br>EDFYLVIHIIHQIQSMKTLWELVNKDPKELEEFLMTHKENLMLK<br>DIASPLKTVYKPCFEEYTKKPKLDHNQEIVKYWYNYHIERYWN<br>TPEAKLEFYRKFGQVDLKQPAILAKFASKIKKNKNYKIYLLPQL<br>VVPTYNAEQLESDVAKEILEYTKLMPEERKELLENILAEVDSDII<br>DKSLSEIEVEKIAQELENKIRVRDDKGNSVPISQLNVQKSQLLL<br>WTNYSRKYPVILPYEVPEKFRKIREIPMFIILDSGLLADIQNFATN<br>EFRELVKSMYYSLAKKYNSLAKKARSTNEIGLPFLDFRGKEKVI<br>TEDLNSDKGIIEVVEQVSSFMKGKELGLAFIAARNKLSSEKFEEI<br>KRRLFNLNVISQVVNEDTLKNKRDKYDRNRLDLFVRHNLLFQV<br>LSKLGVKYYVLDYRFNYDYIIGIDVAPMKRSEGYIGGSAVMFDS<br>QGYIRKIVPIKIGEQRGESVDMNEFFKEMVDKFKEFNIKLDNKK<br>ILLLRDGRITNNEEEGLKYISEMFDIEVVTMDVIKNHPVRAFAN<br>MKMYFNLGGAIYLIPHKLKQAKGTPIPIKLAKKRIIKNGKVEKQ<br>SITRQDVLDIFILTRLNYGSISADMRLPAPVHYAHKFANAIRNEW<br>KIKEEFLAEGFLYFV | Wild type |
| *Mf*Ago | MKNLRYKINAYRIKKDYIPKEVYRYRIRSFIENINIYRFVGFYGG | Wild type |

| | | |
|---|---|---|
| | VALNQSEFILPYPVENLVLEYDGKDVKLEHIDTLNLEDIENKDK<br>EKAEKLVRGYLTSIYKLKPILYKILRDVRESKIINDIRVDPIPDFTV<br>KRHNNEYYLVIDFNHTATVLKNLWDFVGRDKLKLEDYIGKKIIF<br>KPNPKKRYTIKSIEKQNKKDIDDIVEHIIEYYKWTEEEIKSTFGEI<br>DYTQPIIHCEGIPYPFAPQFCNIVFTMEDLDENTLKDLQSYWRLP<br>NEIKGNIINQIAKKLRFVENEPIELEFIKFNNTPLIVKDENGKPTKI<br>YTTNRLFRWNYDSKSKLYLPYDIPDIIKNKTLTTFVLIDENLKNV<br>SGKIKRKVYQMFKNYNKIASKTELPKFDFANKWKYFSNNNIRD<br>VIRKIKDEFNEELGFALIIGNRYYENDYYETLKMQLFNLNIISQNI<br>LWENWSKDDNNFMTNNLLIQIMGKLGIKYFALDAKVNYDYIM<br>GLDSGLGAFKSNRVSGCTVIYDSEGKIRRIQPIDVPSPGERIPIHL<br>VVEFLETKTDINMENKNILFLRDGFVQNSEREELKKLSKELNSN<br>IEVISIRKNNKYKVFTSDYGIGSIFGNDGIFLPHKTTFGSNPVKLS<br>TWLRFNSGNEEKLKINESIMQLLYDLTKMNYSALYGEGRNLRIP<br>APIHYADKFVKALGKNWKIDEELLKHGFLYFI | |
| *Fp*Ago | MMLNIFEVDKNRVEVPQDVYLYKVHLKTLEQRKRDMCIAVLR<br>NSFGYLDVNSFVIYSYKEIRDLPRRIKKYCDLEPTGKVKMNEVD<br>ENVRNALVKTFLRSKIRKDIKKLLKKFKKFQQSVGRWTVALDL<br>ERIELVEHNGEILVSFNVKLNISSMINLWDIIERDVNRLKGLCWS<br>PENLNDNRIWFRYIPHLIIEEVEDFEESAKSFILSDVHTGEEFGG<br>WTTEDLKKYPENEYGLSEDAIKKIGNYTQFDSTQPIIKGVTWSG<br>KEYPFLPQHCIPAYNPMLATAEEKKRIEEIKINLKNKKDEIIRKIIE<br>QLPYLKQPDNIEIKKVQETARLRAKFVKVEVTKGKITKTLSQPY<br>EKPVSSTLDLFGWISRIIDGGDGIIEICIPDYIPENLSRIKEIEAFLLI<br>ENDLNENERKVGDKLLNDAIYVYNFVRSVCLRCGINIPYLNYK<br>GNRFYFENSKEGIRDIYKRITTSLSGEIGFALIFGKRDNYEDEEGE<br>DSFDYYNPLKSALFRNNILSQNFDVTNYVRGDGKINKNTIKYAV<br>SNIIYNIFGKLGVKFFVLEEDVPYDYILGIDVGYGEAYTGKVAGC<br>TTVHDSEGRLRNLIPIEKQNYPSKETARIKALLEEIEQKKKIYNID<br>FENKSILILRDGRINKEEINQLMEFSEERNCRITYIEIRKNIVHQFL<br>VNSSQACYVKIGDYYILKAHNPRIGFPRAIKIARKIVIEGDAWRE<br>SSLTEDDILLIYKLTALNYSTIGRDSNLRIPAPIYYADKLVKALKK<br>GWKFDERFLRYGILYFL | Wild type |

**Table S6.** List of nucleic acids sequences for the designed mismatched gDNA directed cleavage on target DNA

| Name | Sequence (5'-3') |
|---|---|
| 45nt DNA target | ATATACTATACAACCTACTACCTCGTATAAATTTTTAAATAAATA-FAM |
| guide_m0 | P-TGAGGTAGTAGGTTGT |
| guide_m1 | P-AGAGGTAGTAGGTTGT |
| guide_m2 | P-TCAGGTAGTAGGTTGT |
| guide_m3 | P-TGTGGTAGTAGGTTGT |
| guide_m4 | P-TGACGTAGTAGGTTGT |
| guide_m5 | P-TGAGCTAGTAGGTTGT |
| guide_m6 | P-TGAGGAAGTAGGTTGT |
| guide_m7 | P-TGAGGTTGTAGGTTGT |
| guide_m8 | P-TGAGGTACTAGGTTGT |
| guide_m9 | P-TGAGGTAGAAGGTTGT |
| guide_m10 | P-TGAGGTAGTTGGTTGT |
| guide_m11 | P-TGAGGTAGTACGTTGT |
| guide_m12 | P-TGAGGTAGTAGCTTGT |
| guide_m13 | P-TGAGGTAGTAGGATGT |
| guide_m14 | P-TGAGGTAGTAGGTAGT |
